# Perinatal fentanyl exposure leads to long-lasting impairments in somatosensory circuit function and behavior

**DOI:** 10.1101/2020.09.19.304352

**Authors:** Jason B. Alipio, Catherine Haga, Megan E. Fox, Keiko Arakawa, Rakshita Balaji, Nathan Cramer, Mary Kay Lobo, Asaf Keller

## Abstract

One consequence of the opioid epidemic are lasting neurodevelopmental sequelae afflicting adolescents exposed to opioids in the womb. A translationally relevant and developmentally accurate preclinical model is needed to understand the behavioral, circuit, network, and molecular abnormalities resulting from this exposure. By employing a novel preclinical model of perinatal fentanyl exposure, our data reveal that fentanyl has several dose-dependent, developmental consequences to somatosensory function and behavior. Newborn male and female mice exhibit signs of withdrawal and sensory-related deficits that extend at least to adolescence. As fentanyl exposure does not affect dams’ health or maternal behavior, these effects result from the direct actions of perinatal fentanyl on the pups’ developing brain. At adolescence, exposed mice exhibit reduced adaptation to sensory stimuli, and a corresponding impairment in primary somatosensory (S1) function. *In vitro* electrophysiology demonstrates a long-lasting reduction in S1 synaptic excitation, evidenced by decreases in release probability, NMDA receptor-mediated postsynaptic currents, and frequency of miniature excitatory postsynaptic currents, as well as increased frequency of miniature inhibitory postsynaptic currents. In contrast, anterior cingulate cortical neurons exhibit an opposite phenotype, with increased synaptic excitation. Consistent with these canghes, electrocorticograms reveal suppressed ketamine-evoked γ oscillations. Morphological analysis of S1 pyramidal neurons indicate reduced dendritic complexity, dendritic length, and soma size. Further, exposed mice exhibited abnormal cortical mRNA expression of key receptors and neuronal growth and development, changes that were consistent with the electrophysiological and morphological changes. These findings demonstrate the lasting sequelae of perinatal fentanyl exposure on sensory processing and function.

## Introduction

Opioid use dependence and addiction have increased to epidemic proportions, leading to substantial financial and societal health burdens in the United States (Ryan, 2018). While opioid use dependence is highest in the U.S., globally, the prevalence of opioid use disorders has increased by 47% from 1990 to 2016, with women representing one-third of the increase (GBD, 2018). In the U.S., more than 20% of pregnant women enrolled in Medicaid are prescribed opioids. In recent years there was a 14-fold increase in the proportion of pregnant women self-reporting opioid use. As a result of these trends, the incidence of infants born to opioid-using mothers has increased >400% between 2000 and 2012 (Haight et al., 2018; Winkelman et al., 2018; Honein et al., 2019). Not only does opioid use increase the risk of miscarriage, premature birth and stillbirth (Whiteman et al., 2014), newborns exposed to opioids *in utero* may develop neonatal opioid withdrawal syndrome (NOWS), lower birth weight, smaller head circumference, and a higher risk of sudden infant death syndrome (Kandall et al., 1976; McPherson et al., 2015). Current pharmacologic treatment options for newborns with NOWS include opioid formulations of morphine, methadone, and buprenorphine (Sutter et al., 2014). While opioid therapy for newborns with NOWS improves acute withdrawal symptoms, such treatment further increases their exposure to these substances. Of particular concern is the fact that neurodevelopmental deficits may be permanent, as some have been found to persist to adolescence and even adulthood (Lee et al., 2020).

There is a higher risk for sensory-related deficits in adolescents who were exposed to opioids prenatally (Ornoy et al., 2001; Kivisto et al., 2015). Perinatal opioid exposure also increases the risk for other disorders that are characterized by sensory processing deficits—such as attention deficit and autism spectrum disorders (Ayres, 1964; Rogers and Ozonoff, 2005; Robertson and Baron-Cohen, 2017; Balasco et al., 2019; Kilroy et al., 2019), which are also increased by perinatal opioid exposure (Ornoy, 2003; Hunt et al., 2008; Rubenstein et al., 2019).

Nearly all studies of perinatal exposure have focused on the naturally occurring opioid, morphine. However, the number of encounters with synthetic opioids, primarily fentanyl, has increased 300% since 2014 (O’Donnell et al., 2017; Jannetto et al., 2019). Fentanyl is 50 to 100 times more potent than morphine and is responsible for the majority of opioid-related overdose deaths (Volpe et al., 2011; Spencer et al., 2019). Understanding the effects that synthetic opioids have on the developmental trajectory of offspring is critically needed to develop effective treatment and prevention plans. To investigate the persistent neurobiological behavioral, circuit, network, and molecular abnormalities following perinatal fentanyl exposure, we developed a mouse model of such exposure, in which pregnant mouse dams are exposed to fentanyl (Alipio et al., 2020). We showed that newborn pups exhibit signs of spontaneous somatic withdrawal. During adolescence, fentanyl exposed mice displayed abnormal affective-like behavior. By adulthood, fentanyl exposed mice exhibited deficits in auditory processing.

In the present study we further validate this model by determining the effects of different doses of fentanyl on maternal care and health. We then use this model to study the consequences of perinatal fentanyl exposure on sensory behaviors during adolescence, as well as the synaptic mechanisms that subserve them. We focus on the primary somatosensory cortex (S1) and anterior cingulate cortex (ACC), as they play key roles in sensory processing and integration (Mountcastle, 1998).

## Materials and Methods

### Animals

All procedures adhered to the *Guide for the Care and Use of Laboratory Animals* and approved by the Institutional Animal Care and Use Committee at the University of Maryland School of Medicine. Male and female C57BL/6J mice were used and bred in our temperature and humidity-controlled vivarium. Separate cohorts of animals were used for each behavioral test to prevent possible crossover effects between tests. When copulatory plugs were identified, we removed the sires, and added fentanyl or vehicle control to the water hydration pouches. Offspring were weaned at postnatal day (PD) 21 and housed 2 to 5 per cage in single-sex groups. Food and water were available ad libitum, and lights were maintained on a 12-hour cycle.

### Statistical Analyses

Statistical tests were conducted using Prism 8 (GraphPad, San Diego, CA) and sample size was determined using G*Power software suite (Heinrich-Heine, Universität Düsseldorf). Detailed statistical data are listed in Table 1. If there was no statistically significant sex difference or exposure/sex interaction, we grouped animals and analyzed according to exposure conditions, per NIH recommendations for studying sex differences. Parametric tests were used when appropriate assumptions were met, otherwise, nonparametric tests were used. Partial η2 were used to calculate effect size, and Cohen’s *d*, Glass’ delta, or Hedges’ *g* were used for two group comparisons, and selected based on whether parametric assumptions were met. All experimenters were blind to treatment conditions throughout data collection, scoring, and analysis.

### Fentanyl citrate

We used 1, 10, or 100 μg/mL fentanyl citrate (calculated as free base) in 2% (w/v) saccharin, or 2% saccharin (vehicle control) in the water hydration pouches, which were replenished weekly until litters were weaned.

### Maternal care behavior

We assessed maternal behavior as previously described (Alipio et al., 2020). We carried out daily observation sessions of pregnant dams (*n* = 9 to 10 dams per group) in the vivarium from PD 1 to 7, between 9:00 and 10:30 AM, and between 2:00 and 3:30 PM. We scan sampled each dam in real time, once a minute for 30 minutes, yielding 30 scans per session; 2 sessions per day for 7 days yielded a total of 420 scans per dam. We used ethological parameters for assessing in and out of nest care-related behaviors, which comprised of: “licking/grooming”, “active nursing”, “passive nursing”, and “nest building”. Additionally, “self-grooming” and “eating/drinking” were taken as measures of dam self-maintenance whereas “pups out of nest” and “climbing/digging” were taken as measures of neglectful behavior.

### Pup retrieval test

On PD 7, we briefly removed the dam from the home cage and disturbed the nesting material, distributing it throughout the cage (*n* = 9 to 11 dams per group, with a minimum of 4 pups in the litter). We then placed two pups in the corners away from the nest end of the home cage. Then, we reintroduced the dam and measured the latency to sniff a pup, retrieve each of the pups, start nest building and crouch over pups. We terminated the test if a dam did not complete the task within 15 minutes, resulting in a latency of 900 seconds for any behaviors not observed during that time frame.

### Spontaneous somatic withdrawal behavior

We tested mice 24 hours after cessation of drug effect on PD 22 (*n* = 8 to 11 mice per group). We first habituated mice to the testing room for 1 hour and scored behaviors in real time in 5 minute time bins for a 15 minute observation period. Spontaneous somatic withdrawal signs were scored using a modified rating scale (Schulteis et al., 1999). Counted signs included escape jumps and wet dog shakes. Escape jumps equaling 2 to 4 were assigned 1 point, 5 to 9 were assigned 2 points, and 10 or more were assigned 3 points. Wet dog shakes equaling 1 to 2 were assigned 2 points, and 3 or more were assigned 4 points. All other presence signs were assigned 2 points and consisted of 10 distinct behaviors, including persistent trembling, abnormal postures, abnormal gait, paw tremors, teeth chattering, penile erection/genital grooming, excessive eye blinking, ptosis (orbital tightening), swallowing movements, and diarrhea. Counted and presence scores were summed from all 3 of the 5 minute time bins to compute the global withdrawal score.

### Sensory threshold and adaptation

To assess tactile sensitivity, we habituated mice (*n* = 10 to 12 mice per group) to an elevated clear plexiglass box with a mesh bottom for 10 minutes. We applied von Frey filaments of increasing forces to the plantar surface of the hind paw. The filament was applied to the same paw throughout the test. We used the up-down method to determine withdrawal threshold (Dixon, 1965; Chaplan et al., 1994; Deuis et al., 2017). To assess sensory adaptation, we applied a von Frey filament above threshold to the plantar surface of the hind paw, opposite to the hindpaw used during tactile sensitivity testing. The filament was applied once every 30 seconds until the animal stopped responding.

### In vitro slice electrophysiology

We used a modified slice collection method (Ting et al., 2014). We anesthetized adolescent mice (PD 40 to 45; see results section for sample size used in each experiment) with ketamine/xylazine, removed their brains, and prepared coronal slices (300-μm-thick) containing the primary somatosensory (S1) or anterior cingulate cortex (ACC). For recordings, we placed slices in a submersion chamber and continually perfused (2 mL/min) with artificial cerebrospinal fluid (ACSF) containing the following (in mM): 119 NaCl, 2.5 KCl, 1.2 NaH_2_PO_4_, 2.4 NaHCO_3_, 12.5 glucose, 2 MgSO_4_·7H_2_O, and 2 CaCl_2_·2H_2_O. Solutions were saturated with carbogen (95% O_2_ and 95% CO_2_) throughout use.

We obtained whole-cell patch-clamp recordings, in voltage-clamp mode (−70 mV), through pipettes containing the following (in mM): 130 cesium methanesulfonate, 10 HEPES, 1 magnesium chloride, 2.5 ATP-Mg, 0.5 EGTA, 0.2 GTP-Tris, 5 QX-314, and 2% biocytin. In S1 and ACC, we recorded from layer 5 neurons. For miniature postsynaptic current recordings, tetrodotoxin (1 μM) was included in the ACSF. All electrically evoked current responses were recorded at 1.5 times threshold, the minimum stimulation required to produce a current response. We positioned the stimulating electrode below the recorded neuron, in layer 6. To isolate excitatory postsynaptic currents (EPSCs), gabazine (1 μM) was included in the ACSF. To isolate inhibitory postsynaptic currents (IPSCs), 6-cyano-7-nitroquinoxaline-2, 3-dione (CNQX; 20 μM) and DL-2-amino-5-phosphonopentanoic acid (APV; 50 μM) was included in the ACSF. The *N*-methyl-D-aspartate receptor (NMDAR)-mediated component of the EPSC was pharmacologically isolated by including CNQX (20 μM) in the ACSF and by voltage-clamping at 40 mV. Impedance of patch electrodes was 4-6 MΩ. Series resistance <40 MΩ was monitored throughout the recording, and recordings were discarded if series resistance changed by >20%. All recordings were obtained at room temperature.

### In vivo electrocorticogram (ECoG) recordings

We anesthetized mice (PD 35; see results section for sample size used in each experiment) with ketamine/xylazine and subcutaneously implanted a radiotelemetry transmitter (ETA-F_10_, Data Sciences International, Minneapolis, MN) with its leads implanted over the dura above S1 (−1.5 mm from bregma) and the cerebellum (−6.4 mm from bregma). Mice were allowed 10 days to recover from surgery. Mice were acclimated to the behavior testing room for 1 hour before ECoG recordings. ECoGs were recorded with Dataquest A.R.T. acquisition system (Data Sciences International) with cortical ECoG recordings referenced to the cerebellum. Baseline 10 minute ECoG recordings were followed by an intraperitoneal injection of vehicle saline or 10 mg/kg ketamine, and 50 min of post-injections recordings. Each mouse was injected with either ketamine or saline on consecutive days, with the order counterbalanced between animals. We analyzed ECoG recordings with custom-written MATLAB scripts (Version 2019a, Mathworks, MA) and the mtspecgramc routine in the Chronux Toolbox (http://chronux.org) (Bokil et al., 2010). Oscillation power in the γ bandwidth (30-80 Hz) was computed in 10 second time bins from spectrograms for each animal and averaged in 10 minute bins.

### Morphology

We filled cells with biocytin (0.1%) during slice electrophysiology recordings and immediately fixed slices in 4% PFA for 24 to 48 hours following recording (see Results for sample size used in each experiment). We washed slices 3 times for 10 minutes each with 1X phosphate buffered solution (PBS). We incubated slices overnight in Streptavidin conjugated to Cy3 (Jackson Immuno; #016-160-084). We washed slices 3 times for 10 minutes each with 1X PBS and mounted the next day. To assess dendritic arbors, Z-stacks were reconstructed using Neurolucida software (MBF Bioscience, Williston, VT). Sholl analysis was performed by counting basal and apical dendrite intersections across 10 μm concentric circles.

### RNA isolation and RT-qPCR

We collected S1 and ACC tissue punches from adolescent (PD 40) mice which were stored at −80° C until processing (*n* = 12 to 14 mice per group). RNA was isolated and extracted from these samples using Trizol (Invitrogen, Carlsbad, CA) and the MicroElute total RNA kit (Omega Bio-tek, Inc, Norcross, GA) with a DNase step. RNA concentrations were measured on a NanoDrop spectrophotometer, and 400□ng cDNA was then synthesized using an iScript cDNA synthesis kit (Bio-Rad Laboratories, Hercules, CA). mRNA expression changes were measured by RT-qPCR with Perfecta SYBR Green FastMix (Quanta, Beverly, MA) and quantified using a CFX384 system (Bio-Rad Laboratories, Hercules, CA) using the ΔΔCT method as described previously (Fox et al., 2020) using GAPDH as a housekeeping gene. Primer sequences can be found in Table 2.

## Results

### Fentanyl exposure has no effect on dam health or maternal care behavior

As we previously described (Alipio et al., 2020), we model perinatal fentanyl exposure by administering fentanyl citrate (0, 1, 10, or 100 μg/mL) in the drinking water of pregnant mouse dams throughout their pregnancy and until their litters were weaned at postnatal day (PD)_21_. Consistent with our previous study—in which we also administered 10 μg/ml fentanyl in the drinking water—we predicted that fentanyl administration would not influence the dam’s general health (Fig 1; *n* = 9 to 10 dams per group). We compared body weight between different concentrations of fentanyl and vehicle control dams throughout pregnancy until weaning (Fig. 1B). There was no interaction between daily weight and fentanyl exposure groups (Two-way RM ANOVA, *F*_(93, 1023)_ = 0.88, *p* = 0.76), nor a main effect of fentanyl exposure on dam weights (Two-way RM ANOVA, *F*_(3,33)_ = 2.05, *p* = 0.12). We also assessed whether fentanyl exposure influences dams’ liquid consumption (Fig. 1C). We found no interaction between daily liquid consumption and fentanyl exposure groups (Two-way RM ANOVA, *F*_(93,1023)_ = 0.87, *p* = 0.79), nor did fentanyl exposure affect liquid consumption between groups across days (Two-way RM ANOVA, *F*_(3,33)_ = 0.82, *p* = 0.49). We observed similar results with dams’ food consumption (Fig. 1D). There was an interaction, with a medium effect size, between daily food consumption and fentanyl exposure (Two-way RM ANOVA, *F*_(93,1023)_ = 1.49, *p* < 0.002, Partial η^2^ = 0.11), however, a post-hoc multiple comparison analysis revealed no differences between exposure groups across each day (Tukey’s post-hoc, *p* > 0.05). These data suggest that there is no difference in daily food consumption or effect of drug exposure between groups. Together, these data suggest that fentanyl exposure during pregnancy, at any of the concentrations tested, does not influence dams’ total body weight, or liquid and food consumption.

**Figure 1.**
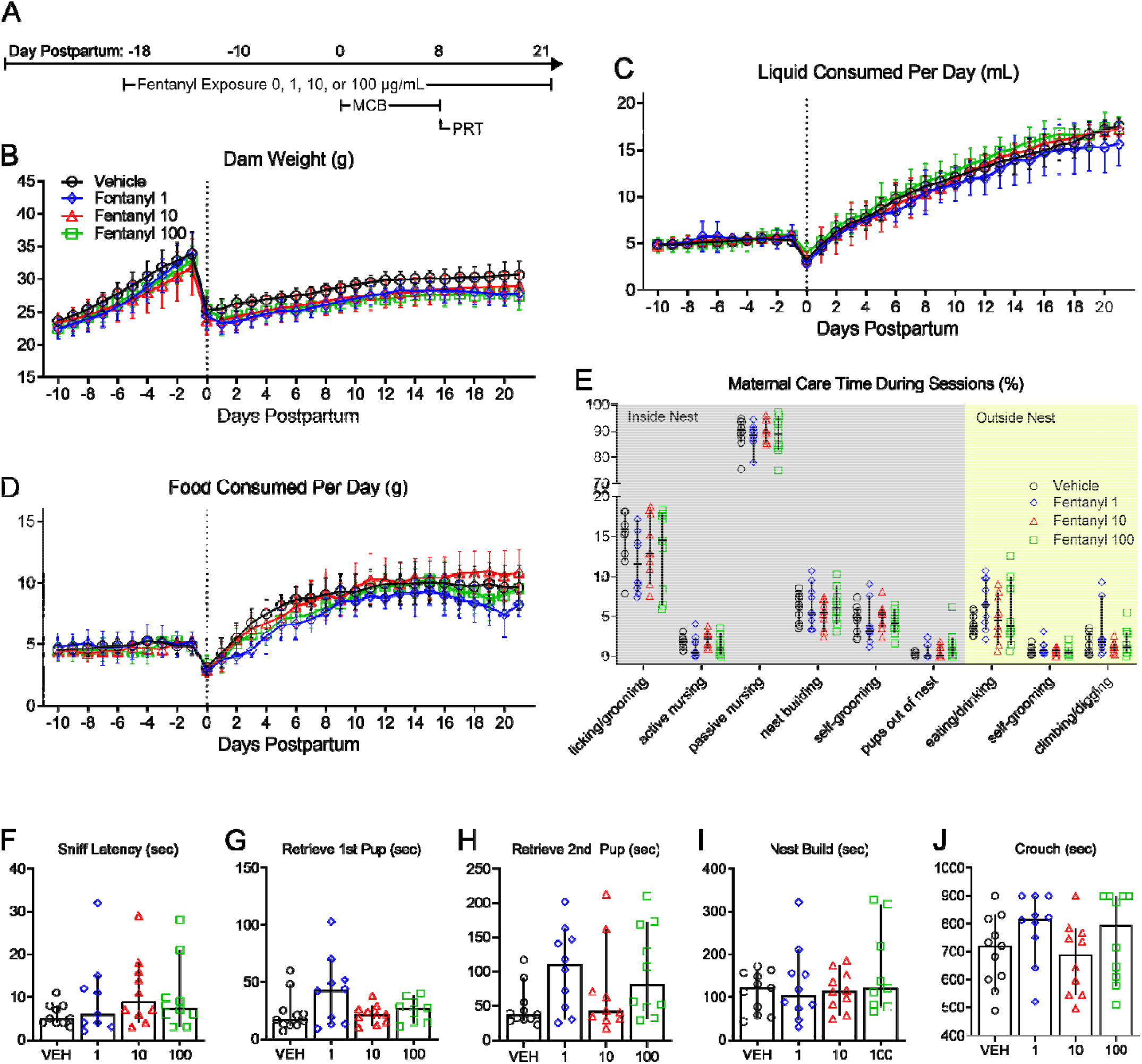
Perinatal fentanyl exposure does not influence dam health or maternal care. **A:** Timeline depicting fentanyl exposure on dams, maternal care behavior, and pup retrieval test. There was no difference in dam weight (**B**), liquid consumption (**C**), or food consumption (**D**) across all concentrations of fentanyl tested. **E:** Fentanyl exposure did not adversely affect passive maternal care behaviors during postnatal days 1 to **F - J**: There were no differences between exposure groups in latencies to sniff, retrieve pups, start nest building, or crouch over pups. Data depict means for parametric or medians for non-parametric comparisons with 95% confidence intervals.

Changes in maternal care during the early postnatal period can influence the development of neural systems, and their associated neurobiological and behavioral outputs (Meaney, 2001; Weaver et al., 2004). To assess maternal care behavior, we compared the time dams spent performing maternal care behaviors during thirty-minute sessions, twice a day, from PD 1 to 7 (Fig. 1E; *n* = 10 dams per group). There was no interaction between fentanyl exposure and maternal care behaviors (Two-way ANOVA, *F*_(24,324)_ = 0.89, *p* = 0.60), nor a main effect of fentanyl exposure (Two-way ANOVA, *F*_(3,324)_ = 0.18, *p* = 0.90), which suggests that fentanyl exposure does not influence maternal care behavior.

To assess a more active form of maternal care, we performed a pup retrieval test on PD 7 (Fig 1F-J; *n* = 9 to 11 dams per group). We compared latencies to sniff a pup, retrieve each of two pups, start nest building, and crouch over pups between fentanyl-exposed and control dams. We found no differences between groups in the latency to sniff a pup (Fig 1H; ANOVA, *F*_(3,35)_ = 1.04, *p* = 0.38), retrieve the first pup (Fig 1G; Kruskal-Wallis test, *H* = 3.26, *p* = 0.35), retrieve the second pup (Fig 1H; ANOVA, *F*_(3,37)_ = 2.36, *p* = 0.08), nest build (Fig 1I; ANOVA, *F*_(3,36)_ = 0.91, *p* = 0.44), or crouch over their pups (Fig 1J; ANOVA, *F*_(3,37)_ = 1.43, *p* = 0.24). Taken together, these data suggest that fentanyl exposure during pregnancy does not adversely affect maternal care behaviors at any of the concentrations tested. We recognize, however, that we did not assess maternal care during the full postnatal period (PD 8 to 21) and cannot exclude possible effects of maternal care during that period.

### Fentanyl exposure results in smaller litters and higher mortality

Women who use opioids during pregnancy have increased incidence of sudden infant death syndrome and miscarriages (Kandall et al., 1975; Kahila et al., 2010; Brogly et al., 2018). We predicted that this increased risk will be observed in litters perinatally exposed to opioids (Fig 2B, C; *n* = 12 litters per group). We compared the number of pups born from dams exposed to different concentrations of fentanyl or vehicle (Fig 2B). There was a large effect of fentanyl exposure on the number of pups per litter (ANOVA, *F*_(3,44)_ = 14.89, *p* < 10^-4^, Partial η^2^ = 0.50), with a large effect decrease in the number of pups per litter at birth that were exposed to each concentration of fentanyl tested, when compared to vehicle controls (Tukey’s post-hoc, 1 μg/mL: *p* < 10^-4^, Cohen’s *d* = 2.51, 10 μg/mL: *p* = 0.001, Cohen’s *d* = 2.39, 100 μg/mL: *p* < 10^-4^, Cohen’s *d* = 2.30). There were no differences in the number of pups per litter between fentanyl exposure groups (*p* > 0.05). We assessed the litter mortality rate at weaning on PD 21 (Fig. 2C). There was a large effect of fentanyl exposure on litter mortality rate (Kruskal-Wallis test, *H* = 13.92, *p* = 0.003, Partial η^2^ = 0.21). Post-hoc analyses indicate a higher mortality rate with large effect sizes for litters that were exposed to each concentration of fentanyl tested, when compared to vehicle controls (Dunn’s post-hoc, 1 μg/mL: *p* = 0.01, Glass’ delta = 3.79, 10 μg/mL: *p* = 0.01, Glass’ delta = 2.92, 100 μg/mL: *p* = 0.01, Glass’ delta = 3.11). There were no differences in litter mortality rate between fentanyl exposure groups (*p* > 0.05). These data indicate that dams exposed to fentanyl during pregnancy, regardless of concentration, have smaller litters and a higher litter mortality rate compared to controls.

**Figure 2.**
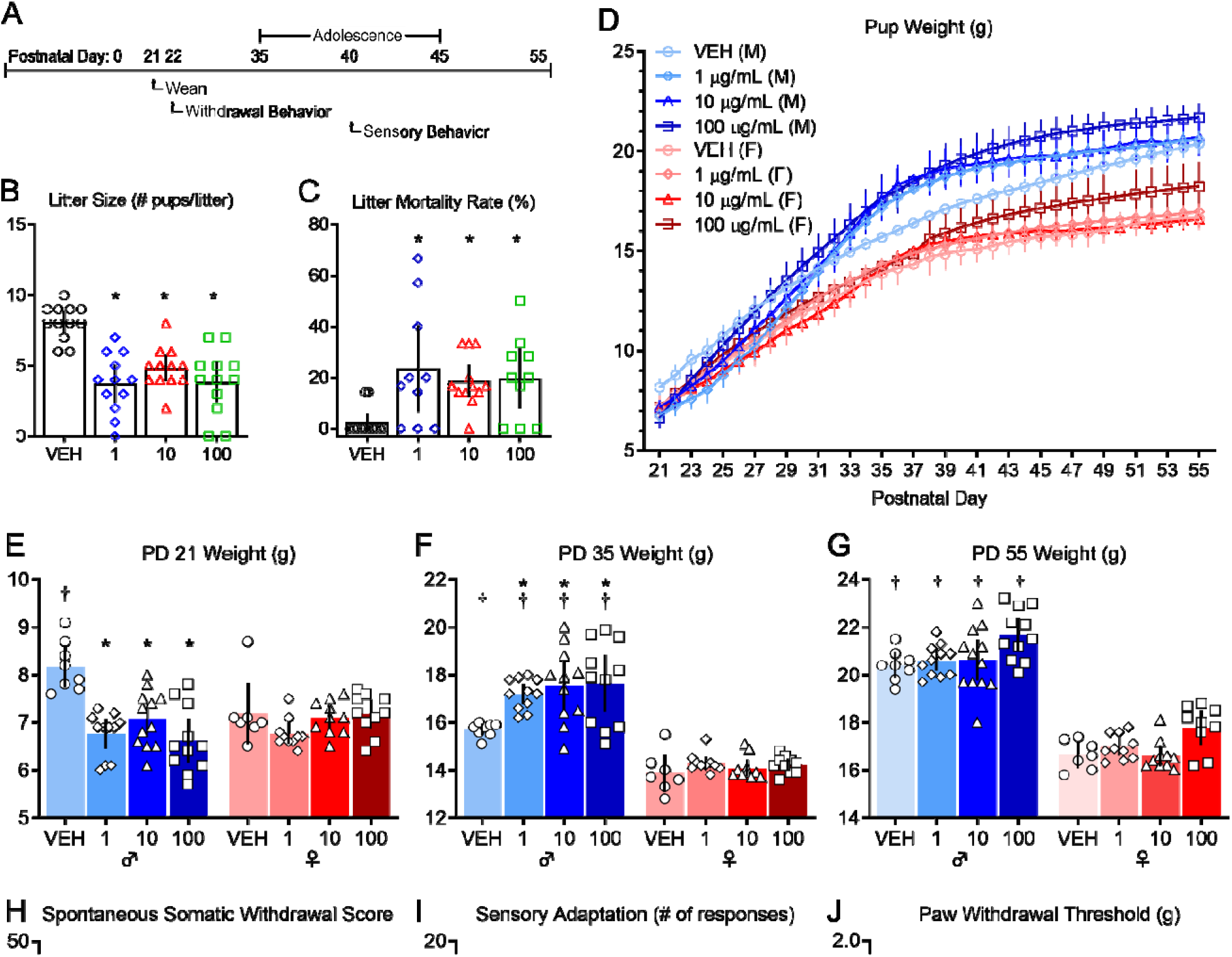
Perinatal fentanyl exposure results in aberrant effects at birth, withdrawal behavior, and impaired sensory function. **A:** Timeline depicting withdrawal behavior 24 hours after weaning and sensory behavior test during adolescence. Exposure resulted in smaller litter size (**B**) and a higher litter mortality rate (**C**). **D**: Mice exposed to fentanyl perinatally exhibited abnormal weight during early development. **E**: At weaning (postnatal day 21), exposed male mice weighed less than controls. **F**: At adolescence (postnatal day 35), exposed male mice weighed more than controls and males weighed more than females at each concentration of fentanyl tested. **G**: By adulthood (postnatal day 55), males weighed more than females, but there were no differences between fentanyl exposure groups. **H**: Perinatal fentanyl exposure induces spontaneous somatic withdrawal behavior 24 hours after cessation, at all concentrations of fentanyl tested. **I**: Perinatal fentanyl exposure impaired sensory adaptation to continuous application of tactile stimuli (**I**), but did not influence paw withdrawal threshold (**J**).

### Abnormal weight across early development

Children prenatally exposed to opioids have lower weights and smaller head circumference compared to age matched controls (Wilson et al., 1979; McPherson et al., 2015). We predicted that pups perinatally exposed to fentanyl would have abnormal body weight across development (PD 21 to 55). To test this prediction, we measured body weight across development of pups perinatally exposed to different concentrations of fentanyl and compared them to controls (Fig. 2D-G; *n* = 8 to 11 mice per group). There was an interaction, with a large effect size, between perinatal fentanyl exposure and weight (Two-way RM ANOVA, *F*_(238,2380)_ = 21.91, *p* < 10^-4^, Partial η^2^ = 0.65), with a large main effect of fentanyl exposure (Two-way RM ANOVA, *F*_(7,70)_ = 30.41, *p* < 10^-4^, Partial η^2^ = 0.83). These data suggest perinatal fentanyl exposure influences body weight across development.

To assess body weight at specific developmental periods, we tested whether pups perinatally exposed to fentanyl had abnormal body weight at weaning (Fig. 2E; PD 21). There was an interaction, with a large effect size, between perinatal fentanyl exposure and sex (Two-way ANOVA, *F*_(3,71)_ = 6.45, *p* = 0.001, Partial η^2^ = 0.23). Exposed males, but not females, weighed less than vehicle controls, with a large effect size (Tukey’s post hoc, 1 μg/mL: *p* < 10^-4^, Cohen’s *d* = 2.82, 10 μg/mL: *p* = 0.004, Cohen’s *d* = 1.96, 100 μg/mL: *p* < 10^-4^, Cohen’s *d* = 2.51). There were no differences in weight between fentanyl exposure groups (*p* > 0.05) and, as expected, vehicle males weighed more than vehicle females (Tukey’s post hoc, *p* = 0.01, Cohen’s *d* = 1.59).

At adolescence (Fig. 2F; PD 35) there was no interaction between perinatal fentanyl exposure and sex (Two-way ANOVA, *F*_(3,70)_ = 2.49, *p* = 0.06). There were main effects of drug exposure (Two-way ANOVA, *F*_(3,70)_ = 4.55, *p* = 0.005, Partial η^2^ = 0.16), and sex (Two-way ANOVA, *F*_(1, 70)_ = 161.2, *p* < 10^-4^, Partial η^2^ = 0.64). Exposed males, but not females, weighed more than vehicle controls, with a large effect size (Tukey’s post hoc, 1 μg/mL: *p* = 0.03, Cohen’s *d* = 2.96, 10 μg/mL: *p* = 0.003, Cohen’s *d* = 1.65, 100 μg/mL: *p* = 0.001, Cohen’s *d* = 1.54). There were no differences in weight between fentanyl exposure groups (*p* > 0.05). Males weighed more than females at all concentrations tested (Tukey’s post hoc, VEH: *p* = 0.01, Cohen’s *d* = 2.82, 1 μg/mL: *p* < 10^-4^, Cohen’s *d* = 5.39, 10 μg/mL: *p* < 10^-4^, Cohen’s *d* = 2.99, 100 μg/mL: *p* < 10^-4^, Cohen’s *d* = 2.69).

By adulthood (Fig. 2G; PD 55) there was no interaction between perinatal fentanyl exposure and sex (Two-way ANOVA, *F*_(3,70)_ = 0.20, *p* = 0.89). There were large main effects of drug exposure (Two-way ANOVA, *F*_(3,70)_ = 7.64, *p* = 0.0002, Partial η^2^ = 0.22), and sex (Two-way ANOVA, *F*_(1,70)_ = 351.9, *p* < 10^-4^, Partial η^2^ = 0.77). In male and female groups, there were no differences in weight when comparing across all fentanyl concentrations tested (Tukey’s post-hoc, *p* > 0.05). Males weighed more than females at all concentrations tested (Tukey’s post-hoc, VEH: *p* < 10^-4^, Cohen’s *d* = 5.94, 1 μg/mL: *p* < 10^-4^, Cohen’s *d* = 6.00, 10 μg/mL: *p* < 10^-4^, Cohen’s *d* = 3.78, 100 μg/mL: *p* < 10^-4^, Cohen’s *d* = 3.83).

Taken together, these results indicate that perinatal fentanyl exposure only affects the body weight of males at weaning. At all other age groups examined, neither males nor females that were treated perinatally differed in weight from age or sex-matched controls.

### Fentanyl withdrawal behavior

Newborns exposed to opioids *in utero* may develop neonatal opioid withdrawal syndrome. Twenty-four hours after weaning (PD 21), at a time when fentanyl was expected to have cleared in treated mice (Hug and Murphy, 1981), we tested whether perinatally exposed mice exhibited spontaneous somatic withdrawal signs (Fig. 2H, I; *n* = 8 to 11 mice per group). There was no interaction between sex and exposure (Two-way ANOVA, *F*_(3,73)_ = 1.81, *p* = 0.15). There were large main effects of sex (Two-way ANOVA, *F*_(1,73)_ = 12.69, *p* = 0.0007, Partial η^2^ = 0.14) and fentanyl exposure (Two-way ANOVA, *F*_(3, 73)_ = 189.5, *p* < 10^-4^, Partial η^2^ = 0.88). Exposed males had higher withdrawal scores compared to controls, with a large effect size, at all concentrations tested (Tukey’s post-hoc, 1 μg/mL: *p* < 10^-4^, Cohen’s *d* = 6.90, 10 μg/mL: *p* < 10^-4^, Cohen’s *d* = 3.29, 100 μg/mL: *p* < 10^-4^, Cohen’s *d* = 8.25). Males exposed to 100 μg/mL had higher withdrawal scores compared to males exposed to 1 μg/mL (Tukey’s post-hoc, *p* < 10^-4^, Cohen’s *d* = 2.08) or 10 μg/mL (Tukey’s post-hoc, *p* < 10^-4^, Cohen’s *d* = 3.46). Males exposed to 1 μg/mL had higher withdrawal scores compared to males exposed to 10 μg/mL (Tukey’s post-hoc, *p* = 0.0009, Cohen’s *d* = 1.75).

Exposed females also had higher withdrawal scores compared to controls, with a large effect size (Tukey’s post-hoc, 1 μg/mL: *p* < 10^-4^, Cohen’s *d* = 4.95, 10 μg/mL: *p* = 0.009, Cohen’s *d* = 2.47, 100 μg/mL *p* < 10^-4^, Cohen’s *d* = 7.00). Females exposed to 100 μg/mL had higher withdrawal scores compared to females exposed to 1 μg/mL (Tukey’s post-hoc, *p* < 10^-4^, Cohen’s *d* = 1.99) or 10 μg/mL (Tukey’s post-hoc, *p* < 10^-4^, Cohen’s *d* = 4.53). Males exposed to 10 μg/mL had higher withdrawal scores compared to females exposed to 10 μg/mL (Tukey’s post-hoc, *p* = 0.01, Cohen’s *d* = 1.42). The expression of withdrawal signs supports the conclusion that our model recapitulates opioid exposure during pregnancy in humans.

### Impaired sensory adaptation in adolescent mice

Adolescent children prenatally exposed to opioids may exhibit lasting somatosensory deficits (Kivisto et al., 2015) that may be independent of the expression of NOWS (Bakhireva et al., 2019).

Adaptation, the cessation of a paw withdrawal response, to repeated stimuli is a key measure of sensory processing. We predicted that adolescent mice perinatally exposed to different concentrations of fentanyl would have impaired adaptation to repeated tactile stimuli (Fig. 2I; *n* = 10 to 12 mice per group). We compared the number of paw withdrawal responses to repeated application of a punctate stimulus that evoked withdrawal responses at threshold forces (see Methods). Compared to control mice, fentanyl exposed mice failed to adapt, that is, they continued to respond to the stimuli. There was an interaction, with a medium effect size, between sex and fentanyl exposure (Two-way ANOVA, *F*_(3,78)_ = 2.736, *p* = 0.04, Partial η^2^ = 0.09). Fentanyl exposed males had a higher number of paw withdrawal responses compared to controls, with a large effect size at all concentrations tested (Tukey’s post hoc, 1 μg/mL: *p* = 0.01, Cohen’s *d* = 1.23, 10 μg/mL: *p* < 10^-4^, Cohen’s *d* = 3.06, 100 μg/mL: *p* < 10^-4^, Cohen’s *d* = 4.28). Likewise, exposed females had a higher number of paw withdrawal responses compared to controls, with a large effect size at all concentrations tested (Tukey’s post hoc, 1 μg/mL: *p* = 0.0002, Cohen’s *d* = 2.30, 10 μg/mL: *p* = 0.02, Cohen’s *d* = 1.57, 100 μg/mL: *p* = 0.0007, Cohen’s *d* = 2.80). In male and female groups, there were no differences in weight when comparing across all fentanyl concentrations tested (Tukey’s post-hoc, *p* > 0.05).

The failure to adapt was not due to reduced sensitivity to stimuli, because the responses to punctate stimuli elicited similar responses in both groups. We used von Frey filaments to test paw withdrawal thresholds (Fig. 2J; *n* = 10 to 12 mice per group). There was no interaction between sex and fentanyl exposure (Two-way ANOVA, *F*_(3,75)_ = 0.38, *p* = 0.76). There was a medium main effect of sex (Two-way ANOVA, *F*_(1,75)_ = 11.39, *p* = 0.001, Partial η^2^ = 0.13) and a large effect of fentanyl exposure (Two-way ANOVA, *F*_(1,75)_ = 4.15, *p* = 0.008, Partial η^2^ = 0.14). However, there were no differences in paw withdrawal thresholds within sex or exposure groups compared to controls (Tukey’s post-hoc, *p* > 0.05). Taken together, these findings suggest that perinatal fentanyl exposure impairs adaptation to repeated tactile stimulation but does not influence mice’s ability to sense tactile stimuli.

### Decreased excitatory synaptic transmission in S1

Behavioral deficits in tactile sensory perception and processing are associated with changes in synaptic transmission in primary somatosensory cortex (Fox et al., 2000). Therefore, we predicted that adolescent mice perinatally exposed to different concentrations of fentanyl would have impaired excitatory synaptic transmission in S1 (Fig. 3). We recorded miniature excitatory postsynaptic currents (mEPSCs) in S1 layer 5 neurons from adolescent mice, in the presence of gabazine, a GABA_A_ receptor antagonist (*N* = 4 to 5 mice per group, *n* = 2 to 5 neurons per mouse). Figure 3B depicts mEPSCs recorded from adolescent male and female mice perinatally exposed to different concentrations of fentanyl or vehicle control. We assessed the inter-event intervals, the reciprocal of their frequency, from three-minute recordings (Fig. 3C). There was a rightward shift in the cumulative frequency plot of the inter-event interval in fentanyl exposed mice compared to controls (Kruskal-Wallis test, *H* = 3244, *p* < 10^-4^, Partial η^2^ = 0.50). The cumulative frequency was shifted to the right in fentanyl exposed male mice compared to controls, with a large effect size at all concentrations tested (Dunn’s post hoc, 1 μg/mL: *p* < 10^-4^, Hedges’ *g* = 4.72, 10 μg/mL: *p* < 10^-4^, Hedges’ *g* = 2.23, 100 μg/mL: *p* < 10^-4^, Hedges’ *g* = 5.16). Similarly, there was a rightward shift in the cumulative frequency of the inter-event intervals in fentanyl exposed females compared to controls, with a large effect size (Dunn’s post hoc, 1 μg/mL: *p* < 10^-4^, Hedges’ *g* = 4.51, 10 μg/mL: *p*< 10^-4^, Hedges’ *g* = 1.09, 100 μg/mL: *p* < 10^-4^, Hedges’ *g* = 3.52).

**Figure 3.**
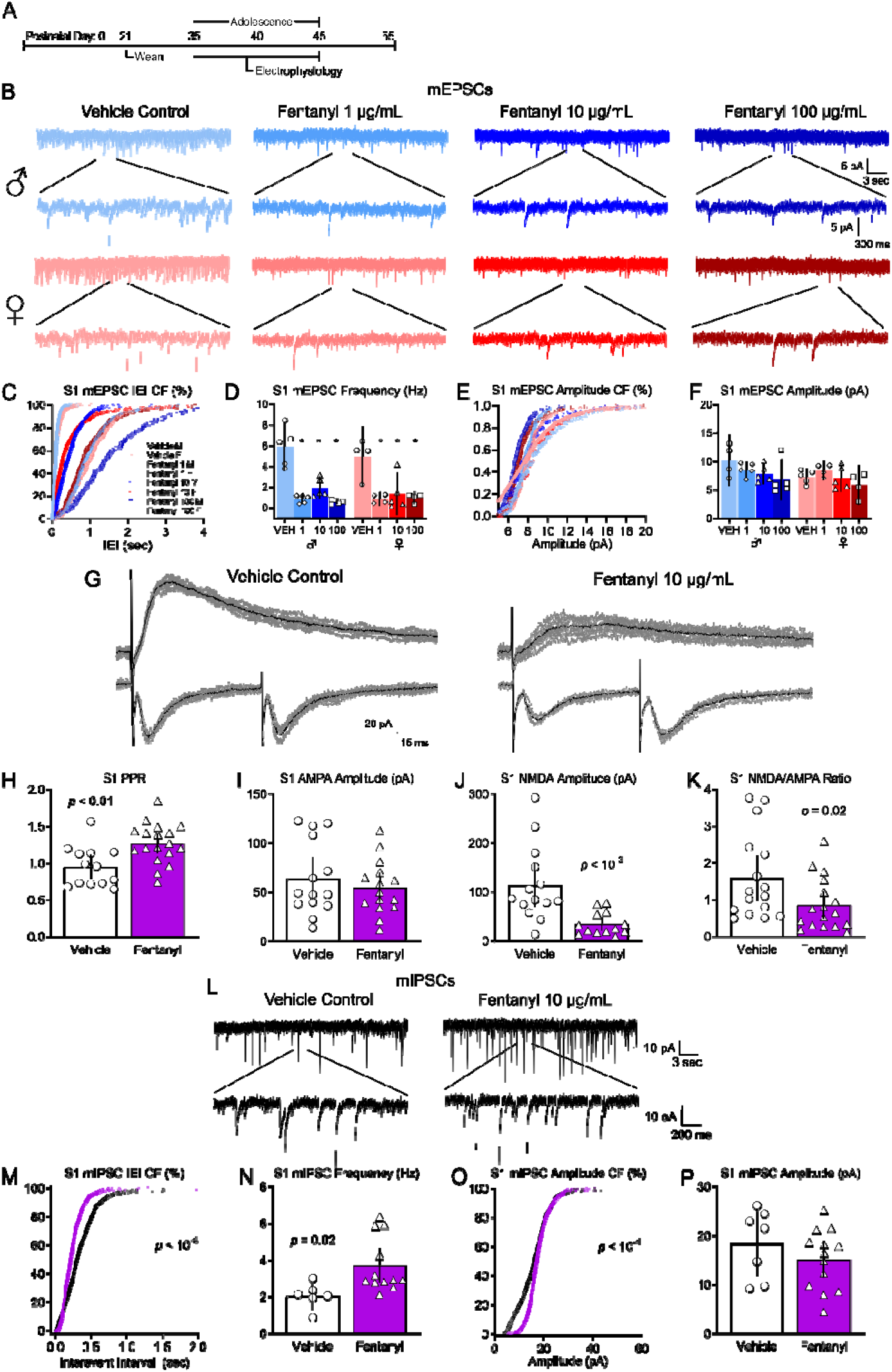
Perinatal fentanyl exposure impairs synaptic transmission in somatosensory cortical neurons. **A:** Timeline depicting slice electrophysiology recordings in S1 layer 5 neurons of adolescent mice. **B:** Example traces of miniature excitatory postsynaptic currents (mEPSCs). **C:** Cumulative frequency plot of the interevent intervals of mEPSCs are shifted to the right in fentanyl exposed mice compared to controls. **D:** Grouped data reflect decreased mEPSC frequency in fentanyl exposed mice. There were no differences in the cumulative frequency of the mEPSC amplitude (**E**) nor in the grouped data for mEPSC amplitude (**F**). **G**: Example traces of evoked paired pulse and NMDA receptor-mediated response. Perinatal fentanyl exposure results in increased paired pulse ratio in fentanyl exposed mice (**H**). There were no differences in AMPA receptor-mediated response amplitude (**I**). **J**: There was decreased NMDA receptor-mediated response amplitude, which is also reflected in the NMDA/AMPA ratio (**K**). **L**: Example traces of miniature inhibitory postsynaptic currents (mIPSCs). **M**: Cumulative frequency plot of the interevent intervals of mIPSCs are shifted to the left in fentanyl exposed mice compared to controls. **N**: Grouped data reflect increased mIPSC frequency in fentanyl exposed mice. **O**: Cumulative frequency plot of the mIPSC amplitude was shifted to the right. **P**: There were no differences in grouped data of mIPSC amplitude.

The frequency of mEPSCs in these samples were lower in fentanyl exposed mice, compared to controls. We averaged data from all neurons recorded from each animal and considered each animal as a single sample (Fig. 3D). This analysis revealed that mEPSCs in fentanyl exposed mice were threefold less frequent than those in controls. There was no interaction between sex and fentanyl exposure (Two-way ANOVA, *F*_(3, 29)_ = 0.57, *p* = 0.63). There was a main effect of fentanyl exposure with a large effect size (Two-way ANOVA, *F*_(3,29)_ = 30.58, *p* < 10^-4^, Partial η^2^ = 0.75) and no effect of sex (Two-way ANOVA, *F*_(1, 29)_ = 0.54, *p* = 0.46). Fentanyl exposed males had a lower frequency of mEPSCs compared to controls, with a large effect size at all concentrations tested (Tukey’s post hoc, 1 μg/mL: *p* < 10^-4^, Cohen’s *d* = 3.66, 10 μg/mL: *p* = 0.0003, Cohen’s *d* = 2.66, 100 μg/mL: *p* < 10^-4^, Cohen’s *d* = 3.93). Similarly, fentanyl exposed females also had a lower frequency of mEPSCs compared to controls, with a large effect size at all concentrations tested (Tukey’s post hoc, 1 μg/mL: *p* = 0.0009, Cohen’s *d* = 3.02, 10 μg/mL: *p* = 0.002, Cohen’s *d* = 2.04, 100 μg/mL: *p* = 0.001, Cohen’s *d* = 2.96). There were no differences in the frequency of mEPSCs between male or female exposure groups regardless of fentanyl concentration (Tukey’s post-hoc, *p* > 0.05) These data indicate that perinatal fentanyl exposure results in lower frequency of mEPSCs in S1 neurons of adolescent mice.

There were shifts in the cumulative frequency of mEPSC amplitudes between fentanyl exposed mice compared to controls, with a small effect size (Fig. 3E; Kruskal-Wallis test, *H* = 543.9, *p* < 10^-4^, Partial η^2^ = 0.21). The cumulative frequency of the amplitude was shifted to the left in fentanyl exposed male mice compared to controls at all concentrations tested (Dunn’s post hoc, 1 μg/mL: *p* < 10^-4^, Hedges’ *g* = 0.35, 10 μg/mL: *p* < 10^-4^, Hedges’ *g* = 0.29, 100 μg/mL: *p* < 10^-4^, Hedges’*g* = 0.57). In female mice, there were no differences between vehicle control and the 1 μg/mL fentanyl group (Dunn’s post hoc, *p* = 0.17), a rightward shift in the 10 μg/mL fentanyl group, with a small effect size (*p* < 10^-4^, Hedges’*g =* 0.24), and a leftward shift in the 100 μg/mL fentanyl group, with a small effect size (*p* = 0.0008, Hedges’ *g* = 0.21).

When data from all cells from a single animal are averaged, and comparisons are made between treated and control animals, we find no differences in amplitudes of mEPSCs in S1 neurons between sex or exposure groups (Fig. 3F). There was no interaction between sex and fentanyl exposure (Two-way ANOVA, *F*_(3,29)_ = 0.78, *p* = 0.51). There was no main effect of fentanyl exposure (Two-way ANOVA, *F*_(1, 29)_ = 2.70, *p* = 0.06) nor a main effect of sex (Two-way ANOVA, *F*_(1, 29)_ = 3.21, *p* = 0.08).

These data suggest that perinatal fentanyl exposure does not impact postsynaptic current responses in adolescent mice.

Since there were no significant interactions between sex and fentanyl exposure, nor a main effect of sex in any of the mEPSC analyses, we grouped and analyzed male and female mice according to exposure conditions. We performed subsequent experiments in mice exposed to 10 μg/mL fentanyl, since there were no robust concentration-dependent difference in the dependent variables tested above. Additionally, this concentration of fentanyl elicits affective and sensory deficits, and is the optimal concentration that mice will readily self-administer (Wade et al., 2008; Alipio et al., 2020).

#### Evoked responses

To determine if fentanyl exposure affects evoked responses, we recorded electrically evoked EPSCs from S1 layer 5 neurons in ACSF containing gabazine to block inhibitory activity. Stimulating electrodes were placed below the recording neuron in layer 6. Figure 3G depicts paired pulse responses recorded from mice perinatally exposed to fentanyl or vehicle control. The paired pulse ratio is calculated by dividing the amplitude of the second pulse by the amplitude of the first pulse. Note that the ratio is higher in the mouse perinatally exposed to fentanyl, compared to the control. Group analysis revealed that perinatal fentanyl exposure led to an increased paired pulse ratio of evoked EPSCs recorded from S1 neurons, which was of a large effect size (Fig. 3H; *N*= 13 to 18 mice per group, *n* = 2 to 5 neurons per mouse, unpaired *t*-test, *t*_(29)_ = 3.14, *p* = 0.003, Cohen’s *d* = 1.14). Because changes in paired pulse ratios primarily reflect changes in presynaptic mechanisms (Zucker and Regehr, 2002), these findings suggest that perinatal fentanyl exposure results in reductions in presynaptic glutamate release in adolescent mice.

#### NMDAR/AMPAR ratios

Figure 3G depicts examples of evoked responses in which the α-amino-3-hydroxy-5-methyl-4-isoxazolepropionic acid receptor (AMPAR) and *N*-methyl-D-aspartate receptor (NMDAR) components of the response were isolated, as described in Methods. Note the smaller NMDA amplitude in fentanyl exposed mouse, compared to the control. Group analysis revealed a reduction in the NMDAR/AMPAR ratio, with a medium effect size (Fig. 3K; *N* = 13 to 17 mice per group, *n* = 2 to 5 neurons per mouse, unpaired *t*-test, *t*_(31)_ = 2.23, *p* = 0.03, Cohen’s *d* = 0.77). This is reflected by a large effect decrease in the NMDAR-mediated EPSC amplitude (Fig. 3J; Mann-Whitney test, *U*= 19, *p* = 0.0002, Glass’ delta = 1.04), and no difference in AMPAR-mediated EPSC amplitude (Fig. 3I; unpaired *t*-test, *t*_(27)_ = 0.79, *p* = 0.43).

Together, these data indicate that perinatal fentanyl exposure impairs spontaneous and evoked excitatory synaptic transmission in S1 neurons in adolescent mice. And that this impairment is mediated both by a reduction in presynaptic glutamate release and a reduction in postsynaptic, NMDAR-mediated responses.

### Increased inhibitory synaptic transmission in S1

Cortical activity involves both excitatory and inhibitory synaptic transmission. To determine if perinatal fentanyl exposure influences inhibitory synaptic transmission in S1, we compared miniature inhibitory postsynaptic currents (mIPSCs) in S1 layer 5 neurons from adolescent mice perinatally exposed to fentanyl or vehicle control (*N* = 7 to 10 mice per group, *n* = 2 to 5 neurons per mouse). Figure 3L depicts mIPSCs recorded from a mouse perinatally exposed to fentanyl or a vehicle control. The frequency of mIPSCs in these samples is higher in the fentanyl exposed mouse. The cumulative probability of the interevent interval was shifted to the left in fentanyl exposed mice compared to controls, with a small effect size (Fig. 3M; Kolmogorov-Smirnov test, *D* = 0.26, *p* < 10^-4^, Hedges’ *g* = 0.20). As a group, fentanyl exposed mice exhibited a large effect increase in mIPSC frequency (Fig. 3N; unpaired *t*-test, *t*_(16)_ = 2.48, *p* = 0.02, Cohen’s *d* = 1.38). These data suggest that perinatal fentanyl exposure increases the frequency of inhibitory spontaneous vesicle release events in S1 neurons from adolescent mice.

There was a rightward shift in the cumulative frequency of mIPSC amplitudes in fentanyl exposed mice, which is of a small effect size (Fig. 3O; Kolmogorov-Smirnov test, *D* = 0.21, *p* < 10^-4^, Hedges’ *g* = 0.29). This difference is driven by a larger number of events at lower amplitudes. However, when data from all cells from a single animal are averaged, and comparisons are made between treated and control animals, we find no difference in mIPSC amplitude (Fig. 3P; unpaired *t*-test, *t*_(18)_ = 1.08, *p* = 0.29).

These data suggest that perinatal fentanyl exposure increases spontaneous inhibitory synaptic transmission in S1 neurons from adolescent mice, and that these changes may be expressed through a presynaptic mechanism.

### Increased excitatory synaptic transmission in ACC

The anterior cingulate cortex (ACC) has been implicated in complex sensory processing behaviors (Kennerley et al., 2006), and behavioral adaptation to environmental stimuli (Brockett et al., 2020). Since perinatal fentanyl exposure impaired sensory adaptation in adolescent mice, we predicted that synaptic transmission in ACC from exposed mice would also be impaired (Fig. 4). Surprisingly, the changes in the adolescent ACC of perinatally treated mice were opposite to those we identified in S1.

**Figure 4.**
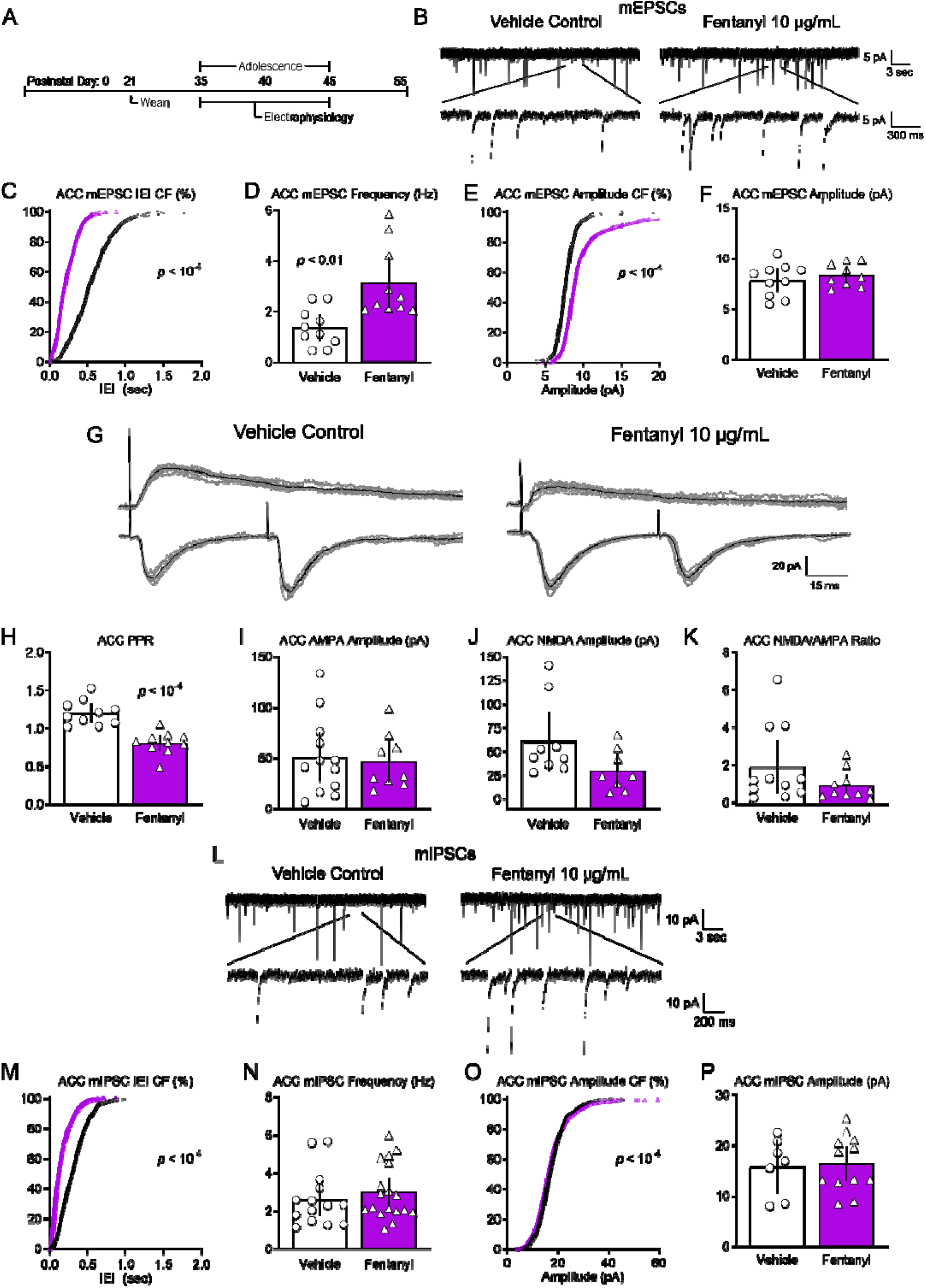
Perinatal fentanyl exposure impairs synaptic transmission in anterior cingulate cortical neurons. **A:** Timeline depicting slice electrophysiology recordings in ACC layer 5 neurons of adolescent mice. **B:** Example traces of miniature excitatory postsynaptic currents (mEPSCs). **C:** Cumulative frequency plot of the interevent intervals of mEPSCs are shifted to the left in fentanyl exposed mice compared to controls. **D:** Grouped data reflect increased mEPSC frequency in fentanyl exposed mice. **E:** Cumulative frequency of the mEPSC amplitude was shifted to the right in fentanyl exposed mice. **F:** There were no differences in grouped data of mEPSC amplitudes **G:** Example traces of evoked paired pulse and NMDA receptor-mediated responses. **H:** Perinatal fentanyl exposure results in decreased paired pulse ratio in fentanyl exposed mice. There were no differences in AMPA receptor-mediated response amplitudes (**I**), NMDA receptor-mediated response amplitudes (**J**), or the NMDA/AMPA ratio (**K**). **L**: Example traces of miniature inhibitory postsynaptic currents (mIPSCs). **M:** Cumulative frequency plot of the interevent intervals of mIPSCs are shifted to the left in fentanyl exposed mice compared to controls. **N:** There were no differences in grouped data of mIPSC frequency. **O:** Cumulative frequency plot of the mIPSC amplitude was shifted to the right. **P:** There were no differences in grouped data of mIPSC amplitude.

We recorded mEPSCs in ACC layer 5 neurons from adolescent mice perinatally exposed to fentanyl or vehicle control (*N* = 8 to 10 mice per group, *n* = 2 to 5 neurons per mouse). Figure 4B includes sample recordings depicting increased frequency and amplitude of mEPSCs recorded in ACC layer 5 neurons from an exposed mouse or a vehicle control. The cumulative frequency plot of the interevent intervals was shifted to the left in fentanyl exposed mice compared to controls, with a large effect size (Fig. 4C; Kolmogorov-Smirnov test, *D* = 0.58, *p* < 10^-4^, Hedges’ *g* = 1.79). As a group, fentanyl exposed mice exhibited a large effect increase of mEPSC frequency in ACC neurons (Fig. 4D; unpaired *t*-test, *t*_(18)_ = 3.43, *p* = 0.003, Cohen’s *d* = 1.53).

The cumulative frequency of the amplitude was shifted to the right in fentanyl exposed mice, with a medium effect size (Fig. 4E; Kolmogorov-Smirnov test, *D* = 0.38, *p* < 10^-4^, Hedges’ *g* = 0.59). However, as a group, there was no difference in the average mEPSC amplitude (Fig. 4F; unpaired *t*-test, *t*_(17)_ = 0.75, *p* = 0.46). As discussed above, this discrepancy might reflect a specific effect on a subset of mEPSCs.

These data suggest that perinatal fentanyl exposure increases the frequency of spontaneous excitatory synaptic transmission in ACC neurons from adolescent mice.

#### Evoked responses

Similarto S1, we investigated if perinatal fentanyl exposure impairs evoked excitatory synaptic transmission in ACC layer 5 neurons. Stimulating electrodes were placed below the recording neuron in layer 6. Figure 4G depicts the paired pulse responses recorded from the ACC of a mouse perinatally exposed to fentanyl or control. Note the decreased paired pulse ratio in the fentanyl exposed mouse. Group analysis revealed that fentanyl exposed mice exhibit a large effect decrease in the paired pulse ratio of AMPAR-mediated EPSCs recorded from ACC neurons (Fig. 4H; *N* = 9 to 10 mice per group, *n* = 2 to 5 neurons per mouse, unpaired *t*-test, *t*_(18)_ = 5.73, *p* < 10^-4^, Cohen’s *d* = 2.56).

#### NMDAR/AMPAR ratios

Figure 4G depicts example traces of evoked AMPAR and NMDAR-mediated currents. There was no difference in the AMPAR-mediated EPSC amplitude (Fig. 4I; *N* = 8 to 11 mice per group, *n* = 2 to 5 neurons per mouse, unpaired *t*-test, *t*_(19)_ = 0.32, *p* = 0.74), the NMDAR-mediated EPSC amplitude (Fig. 4J; Mann-Whitney test, *U* = 16, *p* = 0.05), or in the NMDAR/AMPAR ratio (Fig. 4K; Mann-Whitney test, *U* = 37, *p* = 0.14). These data suggest that perinatal fentanyl exposure increases evoked excitatory transmission to ACC neurons, likely through presynaptic mechanisms.

### No changes in inhibitory synaptic transmission in ACC

Figure 4L depicts mIPSCs recorded from an adolescent mouse perinatally exposed to fentanyl and a control. There was a leftward shift in the cumulative frequency of the interevent interval of fentanyl exposed mice compared to controls, with a large effect size (Fig. 4M; *N* = 14 to 18 mice per group, *n* = 2 to 5 neurons per mouse, Kolmogorov-Smirnov test, *D* = 0.43, *p* < 10^-4^, Hedges’ *g* = 1.16). There was no difference in the average mIPSC frequency between groups (Fig. 4N; unpaired *t*-test, *t*_(30)_ = 0.68, *p* = 0.49).

Likewise, the cumulative frequency of the amplitude was shifted to the left in fentanyl exposed mice compared to controls, with a small effect size (Fig. 4O; *N* = 7 to 12 mice per group, *n* = 2 to 5 neurons per mouse, Kolmogorov-Smirnov test, *D* = 0.08, *p* < 10^-4^, Hedges’ *g* = 0.08). There was no difference in average mIPSC amplitude between groups (Fig. 4P; unpaired *t*-test, *t*_(17)_ = 0.21, *p* = 0.83). These data suggest that perinatal fentanyl exposure does not influence spontaneous inhibitory synaptic transmission to ACC neurons of adolescent mice.

#### Conclusion

Taken together these findings suggest that, in adolescent mice, the largest effects of perinatal fentanyl exposure is decreased presynaptic release of glutamate in S1, and an increase of this release in ACC.

### Reduced cortical oscillatory response

The changes in synaptic efficacies described above suggest that network activity in mice perinatally exposed to fentanyl may be affected. Using electrocorticography (ECoG), we assessed synchronous network activity from large neuronal populations. We and others have previously shown that reproducible ECoG oscillations—particularly in the γ (30 to 80 Hz) range—can be evoked by injecting a sub-anesthetic dose of ketamine (10 mg/kg), an NMDAR antagonist (Raver et al., 2013; Raver and Keller, 2014). We used this approach to compare γ power change in fentanyl exposed and control mice (Fig. 5; *n* = 7 to 9 mice per group). Figure 5B depicts ECoGs recorded from adolescent mice treated perinatally with either fentanyl or vehicle, along with the spectrograms (Fig. 5C, D) of these recordings. Note that the increase in γ-band activity in the treated mouse is smaller than that in the control. Group analysis revealed no interaction between sex and exposure across all time points (Fig 5E; Two-way RM ANOVA, *F*_(4, 56)_ = 1.41, *p* = 0.24), therefore, mice were grouped according to fentanyl exposure. There was a large main effect of fentanyl exposure (*F*_(1,14)_ = 36.74, *p* < 10^-4^, Partial η^2^ = 0.41). Exposed mice exhibited smaller γ-band activity at both the 10-20 minutes (Bonferroni’s post-hoc, *p* = 0.005) and 20-30 minutes (*p* = 0.01) post ketamine time-points. These data suggest that perinatal fentanyl exposure reduces ketamine-evoked γ band cortical activity in awake adolescent mice.

**Figure 5.**
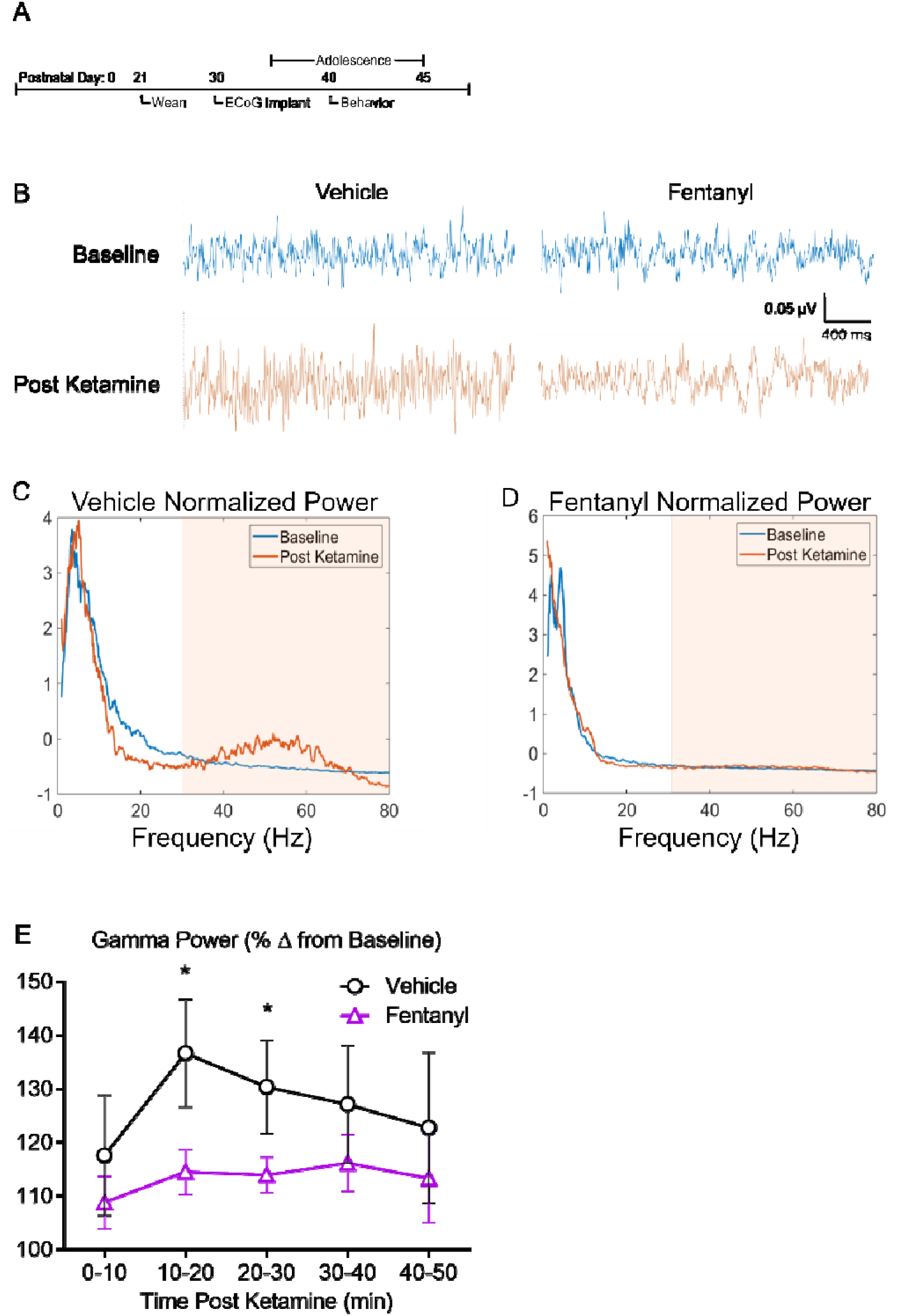
Perinatal fentanyl exposure reduces ketamine-evoked cortical oscillations. **A:** Timeline depicting transmitter implant and *in vivo* electrocorticogram recordings. **B:** Example electrocorticogram (ECoG) traces of cortical oscillations recorded from fentanyl or vehicle control adolescent mice, before and after intraperitoneal injection of sub-anesthetic dose of ketamine (10 μg/mL). Normalized power of cortical oscillations in control **(C)** and perinatal fentanyl exposed **(D)** adolescent mice. **E:** Grouped data of γ power across 10 minute time bins post ketamine injection.

### Reduced branching of pyramidal neurons

Perinatal morphine exposure can delay dendritic development in cortical pyramidal neurons (Ricalde and Hammer, 1990). To determine if perinatal fentanyl exposure influences dendritic morphology we analyzed the morphologies of layer 5 neurons in S1 and ACC (Fig. 6). Note the reduced dendritic complexity in pyramidal neurons from fentanyl exposed mice (Fig. 6B).

**Figure 6.**
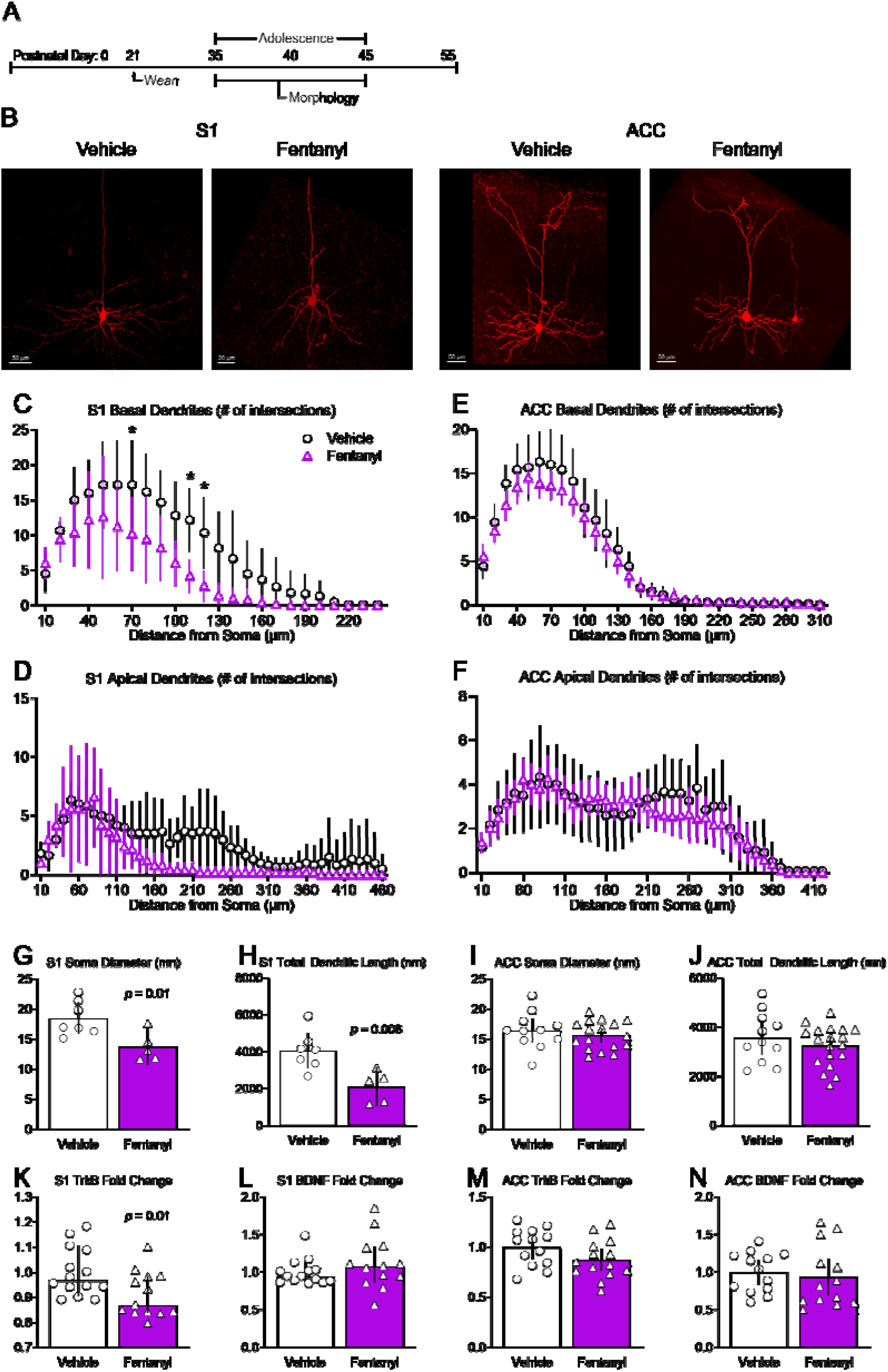
Perinatal fentanyl exposure reduces morphology of basal dendrites of pyramidal neurons in S1. **A:** Timeline depicting morphological assays of pyramidal neurons. **B:** Example of biocytin-filled layer 5 pyramidal neurons in primary somatosensory cortex (S1) and anterior cingulate cortex (ACC). Sholl analysis reveals that perinatal fentanyl exposure results in reduced branching of S1 basal **(C)**, but not apical dendrites **(D)**. There were no differences in branching of ACC basal **(E)** or apical dendrites **(F)**. Fentanyl exposed mice exhibited, in S1, decreased soma diameter **(G)** and decreased total dendritic length **(H)**. Fentanyl exposed mice also had decreased mRNA expression of TrkB receptors K with no difference in expression of BDNF **(L)**. In ACC, there were no differences in soma diameter **(I)**, total dendritic length **(J)**, mRNA expression of TrkB receptors **(M)**, or BDNF **(N)**.

*S1 basal dendrites* from fentanyl exposed mice had less complex branching, evidenced as a smaller number of Sholl intersections (Fig. 6C; *n* = 5 to 7 mice per group). There was a large effect interaction between fentanyl exposure and the number of intersections across dendritic branch distance from the soma (Two-way RM ANOVA, *F*_(23,207)_ = 3.21, *p* < 10^-4^, Partial η^2^ = 0.26), with a smaller number of intersections between 70-120 μm distance from the soma (Bonferroni’s post hoc; *p* < 0.05). Sholl analyses of *S1 apical dendrites* also revealed a main effect of fentanyl exposure (Fig. 6D; Two-way RM ANOVA, *F*_(1,9)_ = 5.99, *p* = 0.03, Partial η^2^ = 0.13), and smaller number of intersections (Two-way RM ANOVA, *F*_(3, 28)_ = 10.89, *p* < 10^-4^, Partial η^2^ = 0.54).

Fentanyl exposure was associated with a large effect reduction in somata diameter (Fig. 6G; unpaired *t*-test, *t*_(10)_ = 2.95, *p* = 0.01, Cohen’s *d* = 1.75), as well as a reduction in total dendritic length (Fig. 6H; unpaired *t*-test, *t*_(10)_ = 3.40, *p* = 0.006, Cohen’s *d* = 2.03) of S1 pyramidal neurons.

As in S1, ACC *basal dendrites* from fentanyl exposed mice had smaller number of Sholl intersections, with a small effect size (Fig. 6E; *n* = 12 to 19 mice per group; Two-way RM ANOVA, *F*_(31,899)_ = 1.50, *p* = 0.03, Partial η^2^ = 0.04). There was no difference in the number of intersections of ACC *apical dendrites* (Fig. 6F). There was no interaction between fentanyl exposure and the number of intersections across dendritic branch distance from the soma (Two-way RM ANOVA, *F*_(41,1189)_ = 0.64, *p* = 0.96), nor a main effect of drug exposure (Two-way RM ANOVA, *F*_(1, 29)_ = 0.17, *p* = 0.67).

There were no differences in somata diameter (Fig. 6I; unpaired *t*-test, *t*_(28)_ = 0.41, *p* = 0.68), nor in total dendritic length (Fig. 6J; unpaired *t*-test, *t*_(28)_ = 0.94, *p* = 0.35) in ACC pyramidal neurons.

Together, these data suggest perinatal fentanyl exposure decreases the dendritic arbor, length, and soma diameter of S1 pyramidal neurons. The dendritic arbor of basal dendrites in ACC were similarly decreased.

### Decreased mRNA expression of TrkB in S1

Brain-derived neurotrophic factor (BDNF) and its receptor, tropomyosin receptor kinase B (TrkB) are involved in growth, development, and maturation of neurons in S1 (Lush et al., 2005). Therefore, the changes in dendritic morphology in pyramidal neurons suggest a corresponding change in the expression of BDNF and TrkB in S1 of adolescent mice (*n* = 13 to 14 mice per group). Indeed, there was a large effect decrease in the expression of TrkB in S1 (Fig. 6K; unpaired *t*-test, *t*_(25)_ = 2.48, *p* = 0.01, Cohen’s *d* = 0.95). There was no difference in the expression of BDNF (Fig. 6L; Mann-Whitney test, *U* = 69, *p* = 0.30). There were no differences in expression of TrkB (Fig. 6M; unpaired *t*-test, *t*_(25)_ = 1.74, *p* = 0.09) or BDNF in the ACC (Fig. 6N; unpaired *t*-test, *t*_(24)_ = 0.48, *p* = 0.63). RT-qPCR primers for BDNF target transcript variants 1 to 12 and primers for TrkB target full and short length isoforms.

These results indicate that decreased mRNA expression of TrkB in S1 corroborates with decreased dendritic arbor, length, and soma size of S1 layer 5 pyramidal neurons. As predicted, there were no differences in expression of BDNF or TrkB in the ACC since there were small effect differences in dendritic arbor.

### Increased CB1R mRNA expression in S1, decreased in ACC

Cannabinoid receptor 1 (CB1R) is a G_i/o_ coupled heteroreceptor and are presynaptically expressed on both inhibitory and excitatory neurons in S1 and ACC (Lomazzo et al., 2017; Yeh et al., 2017). Activation of CB1Rs suppresses presynaptic release. We tested if perinatal fentanyl exposure affects mRNA expression of CB1Rs in S1 and ACC of adolescent mice. In S1, there was a large effect increased expression of CB1Rs (Table 1; unpaired *t*-test, t_(25)_ = 2.41, *p* = 0.02, Cohen’s *d* = 0.92), whereas in ACC there was a large effect decreased expression (unpaired *t*-test, t_(25)_ = 2.75, *p* = 0.01, Cohen’s *d* = 1.05).

These findings demonstrate that changes in CB1R expression parallel the changes in presynaptic release occurring after perinatal fentanyl exposure, suggesting that these changes may be causally related.

### Increased GABA_B_ receptor mRNA expression in ACC

A third mechanism of presynaptic transmitter regulation in neocortex involves GABA_B_ receptors. In ACC, where there was an increase in mEPSC frequency, there was a large effect decrease mRNA expression of GABA_B1_ (Table 1; unpaired *t*-test, t_(24)_ = 2.54, *p* = 0.01, Cohen’s *d* = 0.99). There was no difference in expression of GABA_B2_ receptors (unpaired *t*-test, t_(25)_ = 0.28, *p* = 0.77). In S1, where there was a decrease in mEPSC frequency, there was no difference in expression of GABA_B1_ (unpaired *t*-test, t_(25)_ = 0.27, *p* = 0.78), nor in GABA_B2_ (Mann-Whitney test, *U* = 64, *p* = 0.46). Thus, in ACC, but not in S1, changes in GABA_B_ receptor mRNA expression were correlated with the electrophysiological findings.

### mGluR mRNA expression

In addition to CB1R, metabotropic glutamate receptors (mGluRs) regulate synaptic transmission. We compared the mRNA expression of several mGluR subtypes in adolescent mice perinatally exposed to fentanyl, with controls. In S1, there was a large effect decrease in expression of mGluR_8_ (Table 1; Mann-Whitney test, *U* = 42, *p* = 0.01, Glass’ delta = 0.78). There was no difference in expression of the other group 3 mGluRs: mGluR_4_ (unpaired *t*-test, t_(25)_ = 0.53, *p* = 0.59), or mGluR_7_ (unpaired *t*-test, t_(25)_ = 0.31, *p* = 0.75). There were no differences in group 1 mGluRs: mGluR_1_ (unpaired *t*-test, t_(24)_ = 0.46, *p* = 0.13), and mGluR_5_ (unpaired *t*-test, t_(25)_ = 0.14, *p* = 0.88), nor in group 2 mGluRs: mGluR_2_ (unpaired *t*-test, t_(25)_ = 0.13, *p* = 0.89), and mGluR_3_ (unpaired *t*-test, t_(25)_ = 0.77, *p* = 0.44). Group 3 mGluR_8_ receptors inhibit transmitter release and are expressed on both glutamatergic and GABAergic terminals (Shigemoto et al., 1997; Ferraguti et al., 2005). Given the 3-fold decrease of S1 excitatory transmission, the decreased expression of mGluR_8_ was unexpected. These results suggest that decreased S1 excitatory transmission does not involve changes to mGluR_8_ mRNA expression.

In ACC, where we found a large decrease in presynaptic glutamate release, there was a large effect decrease mRNA expression of mGluR_1_ (unpaired *t*-test, t_(25)_ = 5.82, *p* < 10^-4^, Cohen’s *d* = 2.26), but no difference in expression of mGluR_5_ (unpaired *t*-test, t_(25)_ = 1.23, *p* = 0.22). Among the group 2 mGluRs, there was a large effect decrease expression of mGluR_2_ (unpaired *t*-test, t_(25)_ = 2.55, *p* = 0.01, Cohen’s *d* = 0.98), and mGluR_3_ (unpaired *t*-test, t_(25)_ = 2.13, *p* = 0.04, Cohen’s *d* = 0.81). There were no differences in group 3 mGluRs: mGluR_4_ (unpaired *t*-test, t_(25)_ = 1.45, *p* = 0.15), mGluR_7_ (unpaired *t*-test, t_(25)_ = 0.82, *p* = 0.41), or mGluR_8_ (unpaired *t*-test, t_(25)_ = 2.01, *p* = 0.06). The decreases in group 1 and 2 mGluRs is consistent with a reduction in presynaptic inhibition of glutamate release in ACC, and with our finding of increased excitatory transmission in this cortical area.

### AMPAR and NMDAR mRNA expression

In S1, despite an absence of change in the amplitude of AMPAR currents, there was a large effect decrease in expression of GluR_1_ (Table 1; unpaired *t*-test, t_(25)_ = 2.09, *p* = 0.04, Cohen’s *d* = 0.79), and GluR_3_ (Mann-Whitney test, *U* = 48, *p* = 0.03, Glass’ delta = 1.06). There was no difference in expression of GluR_2_ (Mann-Whitney test, *U*= 62, *p* = 0.16), or GluR_4_ (unpaired *t*-test, t_(25)_ = 1.80, *p* = 0.08). We found no change in the amplitude of AMAR currents in ACC. Consistent with this, there was no difference in expression of GluR_1_ (unpaired *t*-test, t_(25)_ = 1.05, *p* = 0.30), GluR_2_ (unpaired *t*-test, t_(25)_ = 0.05, *p* = 0.95), GluR_3_ (unpaired *t*-test, t_(25)_ = 1.96, *p* = 0.06), or GluR_4_ (unpaired *t*-test, t_(25)_ = 0.09, *p* = 0.92) in ACC of mice perinatally exposed to fentanyl compared to controls. That changes in the expression of GluR_1_ & GluR_3_ did not result in changes in AMPA responses in S1 might relate to the complex interactions and functions among these subunits (Diering and Huganir, 2018).

The reduction in NMDAR-mediated currents in S1 neurons suggests that expression of NMDARs were affected by perinatal fentanyl exposure. However, we found no difference in expression of GluN2A (Table 1; unpaired *t*-test, t_(25)_ = 0.73, *p* = 0.46), GluN2B (Mann-Whitney test, *U* = 84, *p* = 0.75), or GluN_2_D (unpaired *t*-test, t_(25)_ = 0.67, *p* = 0.50) in mice perinatally exposed to fentanyl compared to controls. We did find a large effect increase expression of GluN2C (Mann-Whitney test, *U* = 36, *p* = 0.01, Glass’ delta = 2.04), however these subunits are primarily expressed in S1 astrocytes (Palygin et al., 2011; Ravikrishnan et al., 2018).

In ACC, where there was no change in NMDA currents, there was a large effect decreased expression of GluN2A (unpaired *t*-test, t_(25)_ = 2.52, *p* = 0.01, Cohen’s *d* = 0.96), and GluN2D (unpaired *t*-test, t_(24)_ = 6.17, *p* < 10^-4^, Cohen’s *d* = 2.42). There was no difference in expression of GluN2B (unpaired *t*-test, t_(22)_ = 1.72, *p* = 0.09) or GluN2C (unpaired *t*-test, t_(25)_ = 0.83, *p* = 0.41).

These results may reflect the limited sensitivity of our mRNA expression assay. It is important to note that other molecular mechanisms may be affected by perinatal fentanyl exposure including receptor trafficking, expression of the receptor on the membrane, receptor affinity, or post-translational changes in these subunits. Our expression data is consistent with previous reports showing that prenatal morphine exposure alters kinetic properties of NMDARs (Yang et al., 2000, 2003).

## Discussion

We report that perinatal fentanyl exposure results in neurobiological deficits that last at least until adolescence. This exposure suppresses adaptation to sensory stimuli, impairs synaptic transmission in primary somatosensory (S1) and anterior cingulate cortex (ACC), suppresses cortical oscillations, results in abnormal dendritic morphology of cortical pyramidal neurons, and alters mRNA expression of gene targets related to synaptic transmission and dendritic morphology.

### The model

We confirm and expand upon our previous description of a model of perinatal fentanyl exposure, in which dams are administered fentanyl in their drinking water (Alipio et al., 2020). The treatment extended from conception through weaning, to mimic the embryonic development period in humans (Chen et al., 2017). We expand upon our previous findings by testing the effects of a range of fentanyl concentrations. Here, we find that this exposure has no effect on dam’s health or maternal care behaviors, suggesting that the lasting effects we observe in perinatal fentanyl exposed adolescents reflects the direct actions of fentanyl during early development. Consistent with findings in humans, perinatal fentanyl exposure in mice results in smaller litters, higher newborn mortality rates, and signs of withdrawal shortly after birth. Thus, this model of perinatal opioid exposure has face validity for neonatal opioid withdrawal syndrome and forthe lasting effects of this exposure.

### Cortical excitatory tone

In S1 layer 5 neurons of adolescent mice, perinatal fentanyl exposure resulted in a reduction in synaptic excitation, and an increase in synaptic inhibition. The frequency of miniature excitatory postsynaptic currents (mEPSCs) was increased and that of miniature inhibitory postsynaptic currents (mIPSCs) were decreased, consistent with changes in presynaptic release (Zucker and Regehr, 2002). The increase in the paired-pulse ratio of evoked EPSCs is consistent with reduced vesicle release from excitatory synapses. The amplitude of evoked excitatory synapses was decreased, due, in part, to a reduction in their N-methyl-D-aspartate receptor (NMDAR)-mediated component.

In ACC layer 5 neurons of adolescent mice, perinatal fentanyl exposure enhanced excitatory synaptic transmission. Spontaneous excitatory vesicle release was enhanced, likely through a presynaptic mechanism. This presynaptic effect was mirrored in electrically evoked excitatory synaptic transmission. These findings are demonstrated by increases in the frequency of mEPSCs and depression of evoked paired pulse EPSCs.

Our findings are consistent with previous reports of long-lasting changes in neocortical and hippocampal synaptic activity and plasticity after prenatal exposure to morphine or heroin (Slotkin et al., 2003; Velísek et al., 2003; Yanai et al., 2003; Niu et al., 2009; Tan et al., 2015).

The changes in synaptic transmission in S1 and ACC were large and affected both male and female mice. For example, the frequency of mEPSC were threefold smaller in S1 and twofold larger in ACC than in controls. The NMDA/AMPA ratio was two-fold smaller in S1. These findings indicate that the changes resulting from perinatal fentanyl exposure are physiologically salient.

The contrasting changes in excitatory synaptic transmission in S1 and ACC may be due to differential expression of opioid receptors during cortical development. Opioid receptors are detectable in the central nervous system of mice as early as embryonic day (ED) 11.5 and develop in the cortical subplate at ED 15.5 (Zhu et al., 1998). This is around the same time of migration and differentiation of neurons in layer IV of S1 (Angevine and Sidman, 1961). ACC development does not begin until the last few days of gestation, with full lamination occurring around postnatal day 10 (Van Eden and Uylings, 1985). Activity at these opioid receptors during this critical period of development constrains growth and development of cortical neurons (Zagon and McLaughlin, 1987; Hammer et al., 1989; Ricalde and Hammer, 1990). Thus, the contrasting effects of perinatal fentanyl exposure on S1 and ACC might reflect the different maturational state of opioid systems in these areas at the time of fentanyl exposure.

### Morphological changes

The changes in synaptic functions were accompanied by abnormalities in basal dendrites of S1 layer 5 pyramidal neurons. Consistent with this finding, there was a decrease in mRNA expression of tropomyosin receptor kinase B (TrkB), which is involved in neuronal growth, development, and maturation (Klein et al., 1989). These findings are consistent with prior studies reporting that perinatal morphine exposure reduces dendritic growth and development of S1 pyramidal neurons by acting on opioid receptors during early development (Ricalde and Hammer, 1990; Maharajan et al., 2000; Mei et al., 2009). More recent studies reveal that chronic morphine administration reduces TrkB mRNA expression in D1-type medium spiny neurons within the nucleus accumbens (Koo et al., 2014).

### Neural oscillations

Cortical oscillations, particularly in the gamma range, depend critically on the balance of inhibitory and excitatory synaptic activity (Buzsáki and Wang, 2012). Thus, we predicted that cortical oscillations will become disorganized by perinatal fentanyl exposure. Indeed, treated mice exhibited suppressed oscillations. These oscillations are related to large scale brain network activity and to cognitive phenomena, such as sensory processing, working memory, and attention (Lutz et al., 2004; Steriade, 2006; Buzsáki and Tingley, 2018). Perinatal exposure resulted also in decreased expression of the cannabinoid receptor 1 (CB1R) mRNA, consistent with reports of such decreases after morphine exposure (Rubino et al., 1997). A balance of excitatory and inhibitory transmission is reflected in γ oscillations, that are regulated by CB1R activity (Holderith et al., 2011; Buzsáki and Wang, 2012), suggesting that these synaptic anomalies may be causally related to the oscillation abnormalities. The important role γ oscillations in normal sensory processing and other cognitive tasks suggest that the abnormalities we observed may lead to the lasting deficits in sensory adaptation in mice treated perinatally with fentanyl (Mably and Colgin, 2018; Adaikkan and Tsai, 2020)

### Sensory adaptation

A consequence of the constellation of circuit, network, and morphological effects, reported here, of perinatal fentanyl exposure is compromised sensory adaptation. Here we show that this exposure results in compromised adaptation to repeated tactile stimuli. Longitudinal clinical studies of children and adolescents exposed perinatally to opioids report cognitive, behavioral, and sensory deficits, as well as increased vulnerability to future stressors (Logan et al., 2013; Kivisto et al., 2015; Maguire et al., 2016; Tobon et al., 2019). Impaired sensory processing is associated with attention deficit disorders, autism spectrum, schizophrenia, and synesthesia (Freedman et al., 1987; Lou et al., 1989; Leekam et al., 2007; van Leeuwen et al., 2020).

The synaptic and circuit sequelae of perinatal fentanyl exposure reported here suggest that these lasting behavioral deficits reflect specific and contrasting lasting changes in different cortical regions. Further studies of these changes may direct interventions to ameliorate or prevent these neuropsychiatric deficits.

## Acknowledgements

We thank L. M. Riggs for providing valuable comments on the manuscript. This work was supported by the Opioid Use Disorders Initiative, MPowering The State, from the State of Maryland. Support was provided also by R01 DA038613 to MKL and K99 DA050575 to MEF. The content is solely the responsibility of the authors and does not necessarily represent the official views of the National Institutes of Health. The funding sources had no role in study design; the collection, analysis and interpretation of data; the writing of the report; or in the decision to submit the article for publication.

