## Supplementary material for "Perinatal fentanyl exposure leads to long-lasting impairments in somatosensory circuit function and behavior": Table 1

**Table 1** mRNA expression of receptors involved in synaptic transmission.

| | | Vehicle | | | | Fentanyl | | | | | | |
|---|---|---|---|---|---|---|---|---|---|---|---|---|
| **mRNA** | **Region** | ***n*** | **Mean** | **Median** | **95% CI** | ***n*** | **Mean** | **Median** | **95% CI** | **Analysis** | ***p* value** | ***p* < 0.05** |
| CB1R | S1 | 14 | 1.00 | 0.99 | 0.92-1.07 | 13 | 1.15 | 1.14 | 1.03-1.27 | *t* test | 0.02 | Yes |
| | ACC | 13 | 1.00 | 0.96 | 0.91-1.08 | 14 | 0.86 | 0.85 | 0.78-0.93 | *t* test | 0.01 | Yes |
| $GABA_{B1}$ | S1 | 14 | 1.00 | 1.01 | 0.91-1.08 | 13 | 1.01 | 1.05 | 0.89-1.14 | *t* test | 0.78 | No |
| | ACC | 13 | 1.00 | 0.98 | 0.95-1.04 | 13 | 0.90 | 0.94 | 0.84-0.97 | *t* test | 0.01 | Yes |
| $GABA_{B2}$ | S1 | 13 | 1.00 | 0.97 | 0.89-1.10 | 12 | 1.04 | 1.07 | 0.82-1.27 | Mann-Whitney | 0.46 | No |
| | ACC | 13 | 1.00 | 0.95 | 0.91-1.08 | 14 | 1.02 | 1.01 | 0.85-1.19 | *t* test | 0.77 | No |
| $mGluR_1$ | S1 | 13 | 1.00 | 1.08 | 0.82-1.17 | 13 | 0.93 | 0.93 | 0.66-1.19 | *t* test | 0.64 | No |
| | ACC | 13 | 1.00 | 0.99 | 0.94-1.05 | 14 | 0.73 | 0.70 | 0.65-0.81 | *t* test | $< 10^{-4}$ | Yes |
| $mGluR_2$ | S1 | 14 | 1.00 | 1.06 | 0.88-1.12 | 13 | 1.01 | 0.94 | 0.87-1.15 | *t* test | 0.89 | No |
| | ACC | 13 | 1.00 | 1.05 | 0.91-1.08 | 14 | 0.84 | 0.83 | 0.75-0.94 | *t* test | 0.01 | Yes |
| $mGluR_3$ | S1 | 14 | 1.00 | 0.99 | 0.93-1.06 | 13 | 1.04 | 1.02 | 0.93-1.16 | *t* test | 0.44 | No |
| | ACC | 13 | 1.00 | 0.98 | 0.91-1.08 | 14 | 0.89 | 0.88 | 0.83-0.95 | *t* test | 0.04 | Yes |
| $mGluR_4$ | S1 | 14 | 1.00 | 0.89 | 0.83-1.16 | 13 | 0.94 | 0.85 | 0.80-1.09 | *t* test | 0.59 | No |
| | ACC | 13 | 1.00 | 1.02 | 0.90-1.09 | 14 | 0.90 | 0.91 | 0.80-1.00 | *t* test | 0.15 | No |
| $mGluR_5$ | S1 | 14 | 1.00 | 0.98 | 0.88-1.11 | 13 | 1.01 | 0.95 | 0.86-1.16 | *t* test | 0.88 | No |
| | ACC | 13 | 1.00 | 1.04 | 0.91-1.08 | 14 | 0.88 | 0.79 | 0.70-1.06 | *t* test | 0.22 | No |
| $mGluR_7$ | S1 | 14 | 1.00 | 1.06 | 0.79-1.20 | 13 | 1.04 | 1.13 | 0.79-1.29 | *t* test | 0.75 | No |
| | ACC | 13 | 1.00 | 0.99 | 0.82-1.17 | 14 | 1.21 | 0.79 | 0.68-1.74 | *t* test | 0.41 | No |
| $mGluR_8$ | S1 | 14 | 1.00 | 0.90 | 0.81-1.18 | 13 | 0.74 | 0.74 | 0.69-0.80 | Mann-Whitney | 0.01 | Yes |
| | ACC | 13 | 1.00 | 0.91 | 0.85-1.14 | 14 | 0.83 | 0.86 | 0.73-0.94 | *t* test | 0.05 | No |
| $GluR_1$ | S1 | 14 | 1.00 | 0.99 | 0.90-1.09 | 13 | 0.82 | 0.78 | 0.65-0.98 | *t* test | 0.04 | Yes |
| | ACC | 13 | 1.00 | 0.96 | 0.83-1.16 | 14 | 0.90 | 0.90 | 0.78-1.02 | *t* test | 0.30 | No |
| $GluR_2$ | S1 | 14 | 1.00 | 0.98 | 0.94-1.05 | 13 | 0.92 | 0.86 | 0.76-1.08 | Mann-Whitney | 0.16 | No |
| | ACC | 13 | 1.00 | 1.04 | 0.85-1.14 | 14 | 0.99 | 0.93 | 0.85-1.13 | *t* test | 0.95 | No |
| $GluR_3$ | S1 | 14 | 1.00 | 0.99 | 0.91-1.08 | 13 | 0.84 | 0.78 | 0.66-1.02 | Mann-Whitney | 0.03 | Yes |
| | ACC | 13 | 1.00 | 1.01 | 0.86-1.13 | 14 | 0.83 | 0.91 | 0.72-0.95 | *t* test | 0.06 | No |
| $GluR_4$ | S1 | 14 | 1.00 | 0.95 | 0.92-1.07 | 13 | 0.89 | 0.88 | 0.79-1.00 | *t* test | 0.08 | No |
| | ACC | 13 | 1.00 | 0.96 | 0.91-1.09 | 14 | 1.00 | 1.05 | 0.87-1.14 | *t* test | 0.92 | No |
| GluN2A | S1 | 14 | 1.00 | 1.03 | 0.89-1.10 | 13 | 0.94 | 0.99 | 0.84-1.05 | *t* test | 0.46 | No |
| | ACC | 13 | 1.00 | 1.01 | 0.91-1.08 | 14 | 0.87 | 0.91 | 0.81-0.93 | *t* test | 0.01 | Yes |
| GluN2B | S1 | 14 | 1.00 | 1.00 | 0.88-1.11 | 13 | 1.03 | 0.94 | 0.78-1.28 | Mann-Whitney | 0.75 | No |
| | ACC | 12 | 1.00 | 0.92 | 0.86-1.07 | 12 | 0.86 | 0.83 | 0.77-0.94 | *t* test | 0.09 | No |
| GluN2C | S1 | 14 | 1.00 | 1.00 | 0.91-1.08 | 12 | 1.28 | 1.34 | 1.09-1.48 | Mann-Whitney | 0.01 | Yes |
| | ACC | 13 | 1.00 | 1.08 | 0.86-1.13 | 14 | 0.90 | 0.81 | 0.72-1.09 | *t* test | 0.41 | No |
| GluN2D | S1 | 14 | 1.00 | 0.99 | 0.85-1.14 | 13 | 0.93 | 0.93 | 0.80-1.06 | *t* test | 0.50 | No |
| | ACC | 13 | 1.00 | 1.05 | 0.90-1.09 | 13 | 0.64 | 0.61 | 0.56-0.72 | *t* test | $< 10^{-4}$ | Yes |
