## Supplementary material for "Perinatal fentanyl exposure leads to long-lasting impairments in somatosensory circuit function and behavior": Table 2

**Table 2** RT-qPCR primer sequences.

| Gene | Direction | Sequence |
|---|---|---|
| CB1R | Forward | TGCGCTCTTGCACATGAACT |
| | Reverse | CTGAACGCTGGCCTTACAGA |
| $GABA_{B1}$ | Forward | ACGTCACCTCGGAAGGTT |
| | Reverse | CACAGGCAGGAAATTGATGGC |
| $GABA_{B2}$ | Forward | AAG ACC CCA TAG AGG ACA TCA A |
| | Reverse | GG TGG TAC GTG TCT GTG G |
| $mGluR_1$ | Forward | TGGAACAGAGCATTGAGTTCATC |
| | Reverse | CAATAGGCTTCTTAGTCCTGCC |
| $mGluR_2$ | Forward | GCT CCC ACA GCT ATC ACC G |
| | Reverse | TCA TAA CGG GAC TTG TCG CTC |
| $mGluR_3$ | Forward | CTG GAG GCC ATG TTG TTT GC |
| | Reverse | CAT CCA CTT TAG TCA ACG ATG CT |
| $mGluR_4$ | Forward | CCC ATA CCC ATT GTC AAG TTG G |
| | Reverse | TGT AGC GCA CAA AAG TGA CCA |
| $mGluR_5$ | Forward | CACTGGGGTGCATTGTGTAG |
| | Reverse | GGAGGAGTGCCTGTGTATCA |
| $mGluR_7$ | Forward | AGATGGTGGAATCTTTGGGGA |
| | Reverse | ATTTACAGATGTGCATGGGGG |
| $mGluR_8$ | Forward | CGCTCGCGCAGTGATTATG |
| | Reverse | CCC AAC TAT CTG AGC CAA TCC A |
| $GluR_1$ | Forward | TCC CCA ACA ATA TCC AGA TAG GG |
| | Reverse | AAG CCG CAT GTT CCT GTG ATT |
| $GluR_2$ | Forward | TTC TCC TGT TTT ATG GGG ACT GA |
| | Reverse | CTA CCC GAA ATG CAC TGT ATT CT |
| $GluR_3$ | Forward | ACC ATC AGC ATA GGT GGA CTT |
| | Reverse | ACG TGG TAG TTC AAA TGG AAG G |
| $GluR_4$ | Forward | GTT TTC TGG ATT TTG GGG ACT CG |
| | Reverse | AAG AGA CCA CCT ATT TGA ACG C |
| GluN2A | Forward | ACG TGA CAG AAC GCG AAC TT |
| | Reverse | TCA GTG CGG TTC ATC AAT AAC G |
| GluN2B | Forward | AAAGAGTCGGGGGTGAACTT |
| | Reverse | CGAGGGTTGCCTTCAGTAAG |
| GluN2C | Forward | CATGGCCAGCTTATGACCTT |
| | Reverse | CCAGGACAGGGACACATTTT |
| GluN2D | Forward | GCT GCG AGA CTA TGG CTT CC |
| | Reverse | CCA GTG ACG GGT TTA CCA GAA A |
| TRKB (full and short isoform) | Forward | TTGTGTGGCAGAAAACCTTG |
| | Reverse | ACAGTGAATGGAATGCACCA |
| BDNF (variants 1-12) | Forward | GAAGAGCTGCTGGATGAGGAC |
| | Reverse | TTCAGTTGGCCTTTTGATACC |
| GAPDH | Forward | AGGTCGGTGTGAACGGATTTG |
| | Reverse | TGTAGACCATGTAGTTGAGGTCA |
